# Neural sampling from cognitive maps enables goal-directed imagination and planning

**DOI:** 10.1101/2025.05.14.654027

**Authors:** Hui Lin, Yukun Yang, Rong Zhao, Giovanni Pezzulo, Wolfgang Maass

## Abstract

AI systems are becoming more intelligent, but at a very high cost in terms of energy consumption and training requirements. In contrast, our brains only require 20W of energy, they learn online, and they can instantly adjust to changing contingencies. This begs the question what data structures, algorithms, and learning methods enable brains to achieve that, and whether these can be ported into artificial devices. We are addressing this question for a core feature of intelligence: The capacity to plan and solve problems, including new problems that involve states which were never encountered before. We examine three tools that brains are likely to employ for achieving that: Cognitive maps, stochastic computing, and compositional coding. We integrate these tools into a transparent neural network model, and demonstrate its power for flexible planning and problem solving. Importantly, this approach is suitable for implementation by in-memory computing and other energy-efficient neuromorphic hardware. In particular, it only requires self-supervised local synaptic plasticity that is suited for on-chip learning. Hence a core feature of brain intelligence, the capacity to generate solutions to problems that were never encountered before, does not require deep neural networks or large language models, and can be implemented in energy-efficient edge devices.

## 1 Introduction

Planning and problem solving are at the core of human intelligence. This capability is apparent in multiple domains, such as when planning routes to reach a goal location in a physical environment or when imagining solutions to abstract problems. Importantly, we can apply it also for coping with new challenges that go beyond previous experience.

Numerous experimental data suggest that *cognitive maps*, i.e., data structure for coherently organizing learned experiences by encoding relational information between states and actions, are essential tools that the brain employs for planning. The most widely studied ones are allocentric spatial maps in the rodent hippocampal formation and entorhinal cortex that relate locations in an environment with changes caused by ego-motion (O’Keefe and Nadel, 1978; Behrens et al., 2018; Raju et al., 2024; Moser et al., 2008; George et al., 2021). They rely crucially on the largely innate grid cell system for their formation (Behrens et al., 2018; Hafting et al., 2005; Dong and Fiete, 2024; McNaughton et al., 2006). These cognitive maps support goal-oriented spatial navigation by providing self-localization and a sense of direction towards goals (Epstein et al., 2017; Bush et al., 2015; Tsoar et al., 2011). Recent experimental work in neuroscience and cognitive science goes beyond a purely spatial definition of cognitive maps, suggesting that they are also employed by the brain, especially the human brain, for problem solving in non-spatial domains, including abstract concept spaces (Behrens et al., 2018; Bottini and Doeller, 2020; Tavares et al., 2015; Xiao et al., 2025; Buzsáki and Moser, 2013; Iyer et al., 2024; FeldmanHall et al., 2025).

Multiple lines of evidence suggest that cognitive maps of the brain not only encode past experience but also serve a prospective function, commonly referred to as *imagination*, which enables individuals to anticipate and prepare for situations they have never directly experienced. A rodent study showed that sampling from cognitive maps during replay of sequences of place cells in the hippocampus can be decoded as imagined trajectories in 2D spaces that resemble locomotion plans (Pfeiffer and Foster, 2013; Ólafsdóttir et al., 2018; Vollan et al., 2025). Furthermore, when novel barriers are introduced in the environment, replay trajectories depict goal-directed trajectories around them (Widloski and Foster, 2022).

The human brain is able to imagine and plan also in compositional domains, such as natural language, where a virtually unlimited repertoire of new states and potential goals can be described through new combinations of known components. The underlying brain processes have recently been analyzed in (Schwartenbeck et al., 2023) for composition of novel 2D silhouettes from a set of building blocks (tiles), or more precisely for the NP-hard task to decide whether a given silhouette can be decomposed into these building blocks.

We integrate these three identified tools that brains employ for planning and problem solving, cognitive maps, sampling, and compositional computing, into a novel model for planning and problem solving. This model can be implemented through simple and transparent neural networks that do not require Backprop or Backprop through time for learning. Instead, it learns autonomously in an online manner from own experiences, using only biologically plausible rules for local synaptic plasticity that are easy to implement in neuromorphic hardware. We examine functional capabilities of this model in three very different application domains.

Reinforcement learning (RL) is the standard tool in AI for learning to solve planning or more general problem solving tasks (Sutton and Barto, 2018). However, most RL methods require relearning and substantial computational overhead when the goal changes, and often also deep learning. In contrast, our model can instantly adapt to changes in goals and contingencies through local synaptic plasticity.

Surprisingly, cognitive maps provide a sense of direction even in complex compositional task domains such as composing and decomposing silhouettes from given building blocks. Essential for that is that the standard forward model that defines the relational structure of a cognitive map is complemented by an inverse model that maps state difference to neural codes for actions which cause these state differences. Furthermore, very simple inverse models for cognitive maps often suffice, which provide especially attractive generalization capabilities. In general, inverse models have become a standard modeling tool for biological motor control and in robotics (Wolpert and Kawato, 1998; Kawato, 1999), but they have apparently not yet been considered in the context of cognitive maps.

We will first demonstrate functional abilities of our model in 2D spatial navigation. We use there a simple model for learning cognitive maps that is based on the grid cell system, which uses path integration to update the spatial location that is encoded by grid cell firing (McNaughton et al., 2006; Dong and Fiete, 2024). To address goal-directed sampling in non-spatial domains, we use a simple model for learning cognitive maps that is based on learning predictions of action outcomes through local and biologically-plausible synaptic plasticity rules (Stöckl et al., 2024). We introduce a generative variant of this Cognitive Map Learner (CML) model, the *Generative Cognitive Map Learner (GCML)*, where observations of next states that result from own actions are replaced by internally generated state predictions. If noise is present in the selection of virtual next actions, one arrives at a probabilistic generative model that permits sampling of possible trajectories to any given goal in the cognitive map. Importantly, we show that cognitive maps provide a sense of direction even in complex compositional task domains such as composing and decomposing silhouettes from given building blocks.

## 2 Results

### 2.1 Sampling from a 2D cognitive map reproduces brain data on imagined spatial trajectories

Numerous neural recordings during self-generated imagination processes in the brain are available from the rodent hippocampus. In particular, it was shown in (Pfeiffer and Foster, 2013) that before navigation towards a known goal location (“home”), the hippocampus produced during specific resting phases (short wave ripples), sequential activity of place cells that could be decoded as imagined trajectories from the current location to the goal location. These decoded trajectories exhibited a startling diversity, even for the same starting and goal location, and were rarely straight (see Fig. 2 and Suppl. Fig. 11 and 17 of (Pfeiffer and Foster, 2013)). A subsequent study showed that these decoded trajectories can flexibly reach goals even when this requires rerouting around novel barriers (Widloski and Foster, 2022). These findings cannot be easily accounted for by standard sampling approaches used in statistics and machine learning, such as Markov Chain Monte Carlo (MCMC) and Monte Carlo Tree Search (MCTS) (Bishop, 2006; Gelly and Silver, 2011; Buesing et al., 2011), since these are lacking the goal directed feature. Hence new model for goal-directed stochastic sampling is needed, which we will present here.

We start with a standard model for a cognitive map for spatial navigation that is based upon the grid cell system, see Fig. 1a, and Methods for details. Grid cells produce an allocentric map of a 2D environment based on path integration, thereby relating ego-motion to estimates of spatial locations in the 2D environment (McNaughton et al., 2006; Behrens et al., 2018; Hafting et al., 2005; Liu et al., 2023; Dong and Fiete, 2024). Experimental data suggest that the grid cell system is largely genetically programmed in the rodent, but its development is sped up through spatial navigation (Ulsaker-Janke et al., 2023). Importantly for our context, it has been shown in (Vollan et al., 2025) that the grid cell systems is instrumental in generating imagined trajectories (forward sweeps) in the absence of movements. In fact, their data suggest the presence of a forward model where grid cell states are sequentially updated in the direction of an imagined movement. The allocentric map formed by the grid cell system corresponds on a large scale to a torus (Gardner et al., 2022). We focus on a local segment of the torus, which is approximately flat (Fig. 1b). Whereas each grid cell fires for numerous locations in 2D, they can be used as basis for training place cells that each fire only around a specific location, see Fig. 1a.

**Fig. 1:**
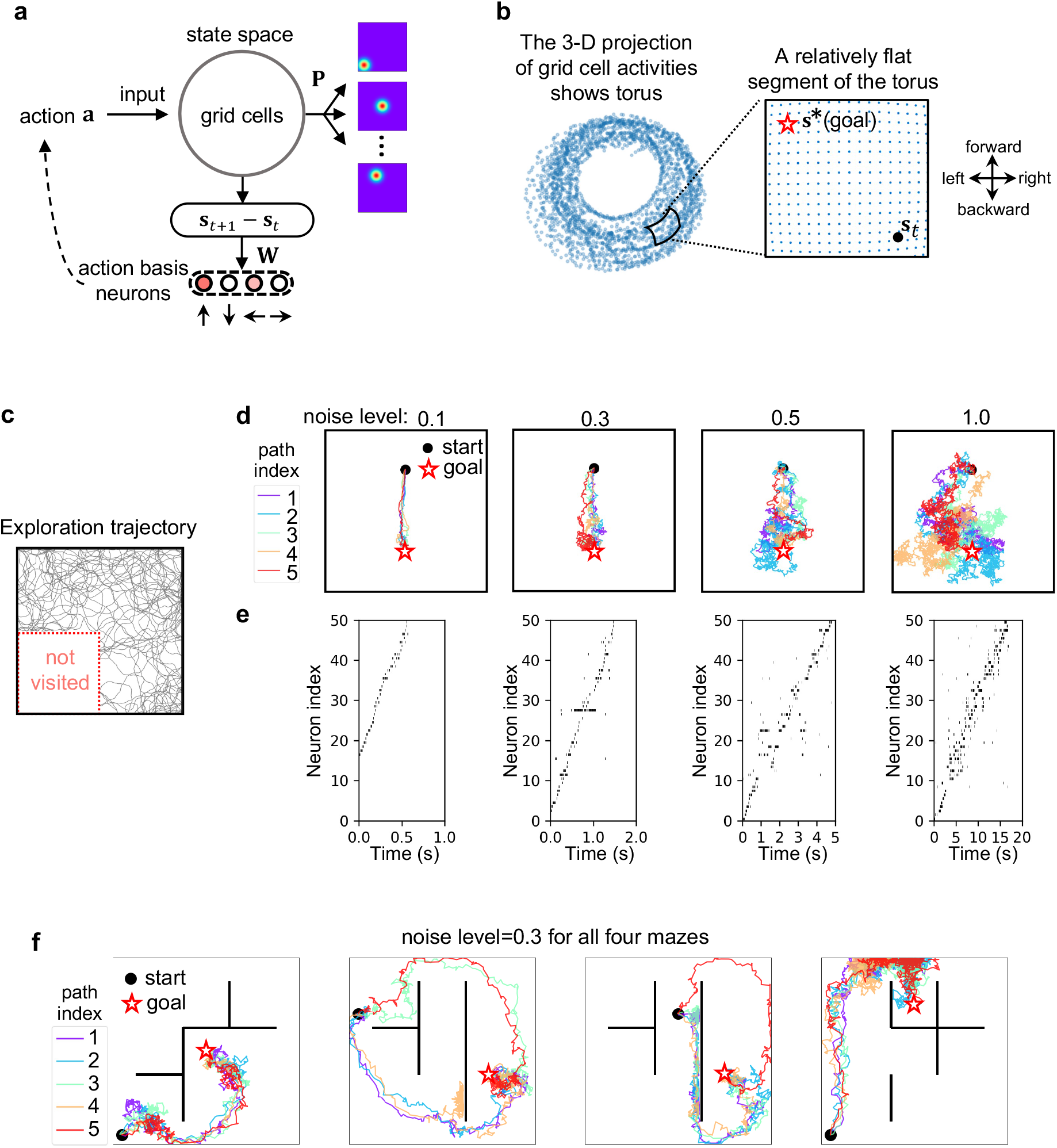
Sampling from a 2D cognitive map supports goal-directed imagination of spatial trajectories. (a) Model architecture. A standard path-integration model for grid cells (Campbell et al., 2018; Sorscher et al., 2023) is used to update grid cell firing on the basis of movement commands (actions) *a*. These are represented here as (normalized) superpositions of basic movement commands in 4 directions. Grid cells provide synaptic inputs to an array of place cells that are trained through a simple local learning mechanism. Simultaneously, also synaptic connections **W** from grid cells to basis action neurons are learned by a simple local synaptic plasticity rule, yielding a linear inverse model. (b) The output of grid cells (for a specific scale) maps a 2D spatial environment onto a segment of the surface of a torus (Gardner et al., 2022). If the grid cells operate on a sufficiently large scale, this segment is approximately flat for a given 2D arena, thereby supporting navigation based on a sense of direction in 2D. (c) Exploration trajectory used to learn the inverse model **W**. We deliberately left a quarter of the space unvisited, where the trajectory is reflected at its perimeter, in order to test whether our model can generate imagined trajectories that also visit unexplored parts of the 2D arena. (d) Impact of different noise levels (0.1, 0.3, 0.5, and 1.0) during action generation on the variance of the generated trajectories. We show five trajectories for each noise level. (e) Spike trains of 50 selected place cells for one of the trajectories (marked in red) in each corresponding plot of panel d. (f) Goal-directed trajectories generated by the same model where repulsive forces from obstacles affect action selection. These trajectories look remarkable similar to those observed in the rodent brain (Widloski and Foster, 2022).

**Fig. 2:**
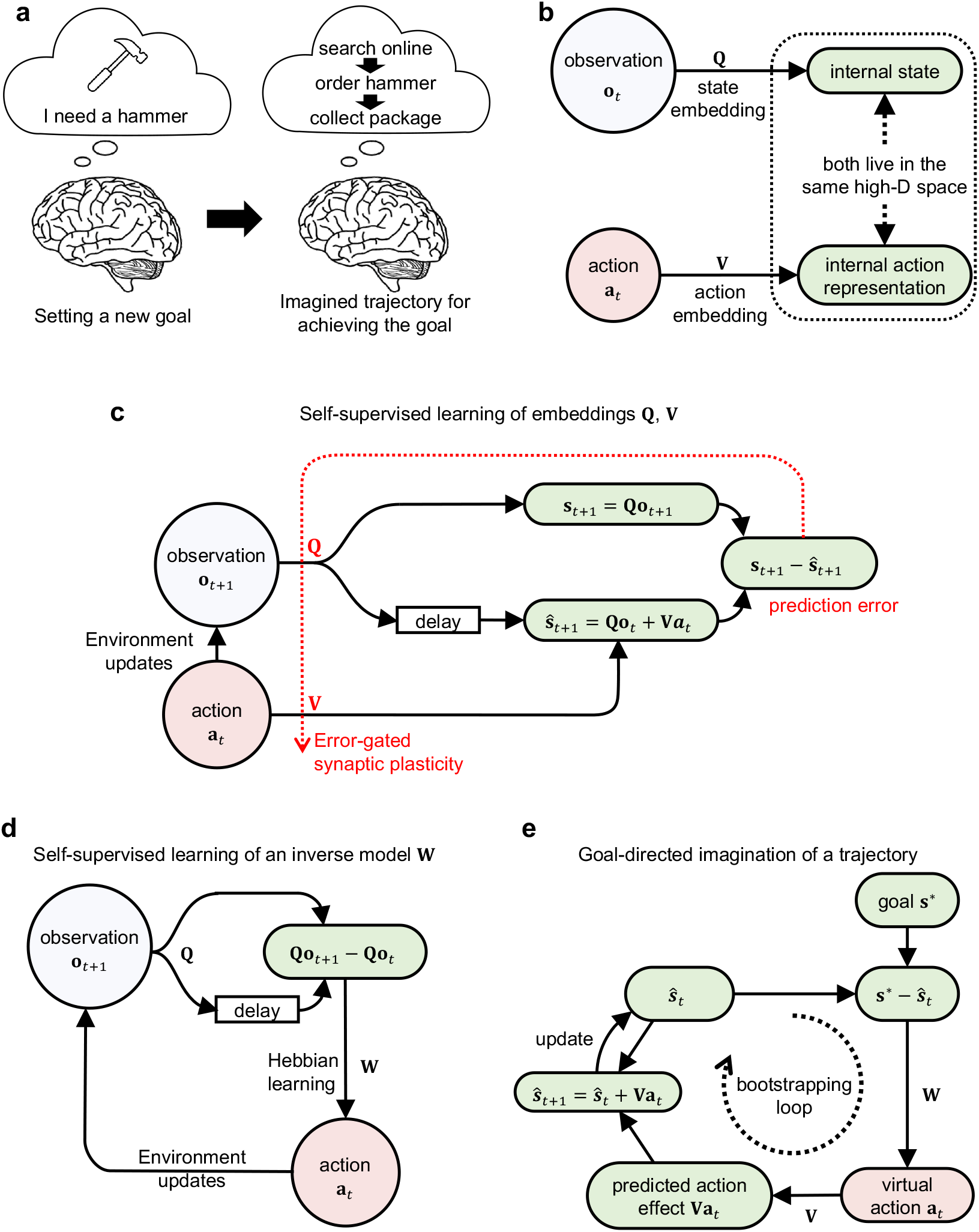
Architecture and learning process of the generative cognitive map learner (GCML). (a) Illustration of a non-spatial problem which brains are able to solve through goaldirected imagination: A plan can be imagined for reaching any given goal, such as getting a hammer. (b) The GCML employs the same cognitive map learning strategy as the CML model of (Stöckl et al., 2024): The cognitive map arises by learning embeddings of observations and (efferent copies of) actions into a common high-D state space so that the geometrical structure of the resulting cognitive map (embeddings) captures the relations between observations and actions. These embeddings **Q** and **V** can be chosen to be linear for the tasks that we consider, and can be learned by simple local rules for synaptic plasticity (c) The learning of the cognitive map, i.e., of the embeddings **Q** and **V**, is driven by a prediction learning goal: Predicting the embedding of the next observation (i.e., the next state) in terms of the current observation and action according to Equ. 11. (d) Simultaneously an inverse model in the form of a matrix **W** can be learnt that maps state differences onto actions which caused them. Note that if the learnt embedding **V** (the forward model) is linear and if Equ. 11 is satisfied, the existence of a linear inverse model is guaranteed. (e) To generate imagined trajectories toward a target state *s*\*, the estimated current state *ŝ*_t_ is iteratively updated. However, importantly, no actions are carried out. Rather, predictions **ŝ**_t+1_ of resulting next observations (states) replace actual feedback from the environment. The next action is virtually selected for the next predicted state (bootstrapping). The difference between a goal state **s**\* and the estimated current state **ŝ**_t_ drives action selection through the learned matrix **W** like for real action selection, thereby enabling goal-directed imagination.

Previous analyses and models suggested that the diversity of 2D trajectories that are decoded from sequential activity of place cells exhibits signs of an underlying stochastic sampling process (Ujfalussy and Orbán, 2022; Stoianov et al., 2022). But a model was missing that could reproduce sampling of goal directed trajectories, such as the virtual trajectories to a “home” location in the rodent (Pfeiffer and Foster, 2013). We show here that such a model emerges if one complements the standard model for the representations by grid- and place cells by a simple self-supervised learning process for an inverse model, that learns during active locomotion to map small changes in grid cell activity to local movement commands which caused them. This inverse model can be implemented by a weight matrix **W**, see Fig. 1a. It is analogous to numerous kinds of inverse models that have previously been employed for modeling biological motor control (Wolpert and Kawato, 1998; Kawato, 1999).

Learning of **W** during exploration of the 2D space (see Fig. 1c) can be implemented by a simple local plasticity rule for synaptic connections from grid cells to action-generating neurons. Due to the approximate linearity of the representation of the 2D space by grid cells and the linearity of **W** one can use this learnt map from state changes to action commands also for selecting a first action that moves into the direction of a distant goal **s**\*, see Fig. 1b and the underlying theory presented in (Stöckl et al., 2024). If this action selection is in addition subject to noise, a model for goal directed stochastic sampling of trajectories emerges. Specifically, the **W** matrix maps the difference between grid cell codes (**s**\*−**ŝ**_t_) for a given goal and the current location onto a superposition (*a*) of basic action commands that would cause, if executed, a movement step from the current location into the direction of the goal. If one replaces the grid cell representation of the next location that would result from this movement step by an estimate that is produced by the forward model, one can iterate this process, yielding an imagined sequence of steps in the direction of the goal. The variance of these imagined trajectories depends on the level of the noise that is superimposed on each action selection, see Fig. 1d. The diversity of trajectories to the given goal that are generated in this way by the model is in the high noise regime qualitatively similar to the one shown in Fig. 4b of (Pfeiffer and Foster, 2013). This predicts that the underlying level of noise in the hippocampus during generation of imagined trajectories is fairly high. Fig. 1e shows the spiking activity of place cells (based on the input which they receive from grid cells according to the architecture of Fig. 1a). More precisely, the underlying spike trains that generate the red-colored trajectories in Fig. 1d are shown. They correspond to recorded spike sequences of place cells in replay studies (Foster, 2017), thereby providing a link to biological data. But place cells or spiking neurons, are not required for the functioning of our model.

Importantly, the goal-directed sampling of our model has an inherent generalization mechanism, allowing the generation of imagined movement trajectories that traverse parts of the space never encountered during training, as observed in the rodent (Gupta et al., 2010). During learning of **W** a large part of the 2D space was not accessible, see Fig. 1c. But yet the sampled trajectories can pass through it, as shown in the plots for high noise levels in Fig. 1d.

Finally, we include in the model the contribution of hippocampal object cells (Manns and Eichenbaum, 2009) and barrier cells (Rivard et al., 2004), whose firing encodes distance and direction of an obstacle or wall. We model the contribution of these cells as repulsive forces from obstacles and walls during the stochastic action selection. As a result, the sampled goal-directed trajectories avoid objects and walls (Fig. 1f). Note that in this simulation, the path requires temporally moving away from the goal vector, to avoid the obstacle, but yet the selected paths maintain a sense of direction towards the goal. This feature becomes even more prominent when larger detours are required, see Fig. S1. This model provides a potential explanation for the puzzling finding that hippocampal replay trajectories show flexible rerouting around novel barriers, without changes (remapping) in place fields (Widloski and Foster, 2022). In fact, the goal-directed trajectories that the model generates look remarkable similar to the recorded ones in that study, see e.g. their Figure 3a.

**Fig. 3:**
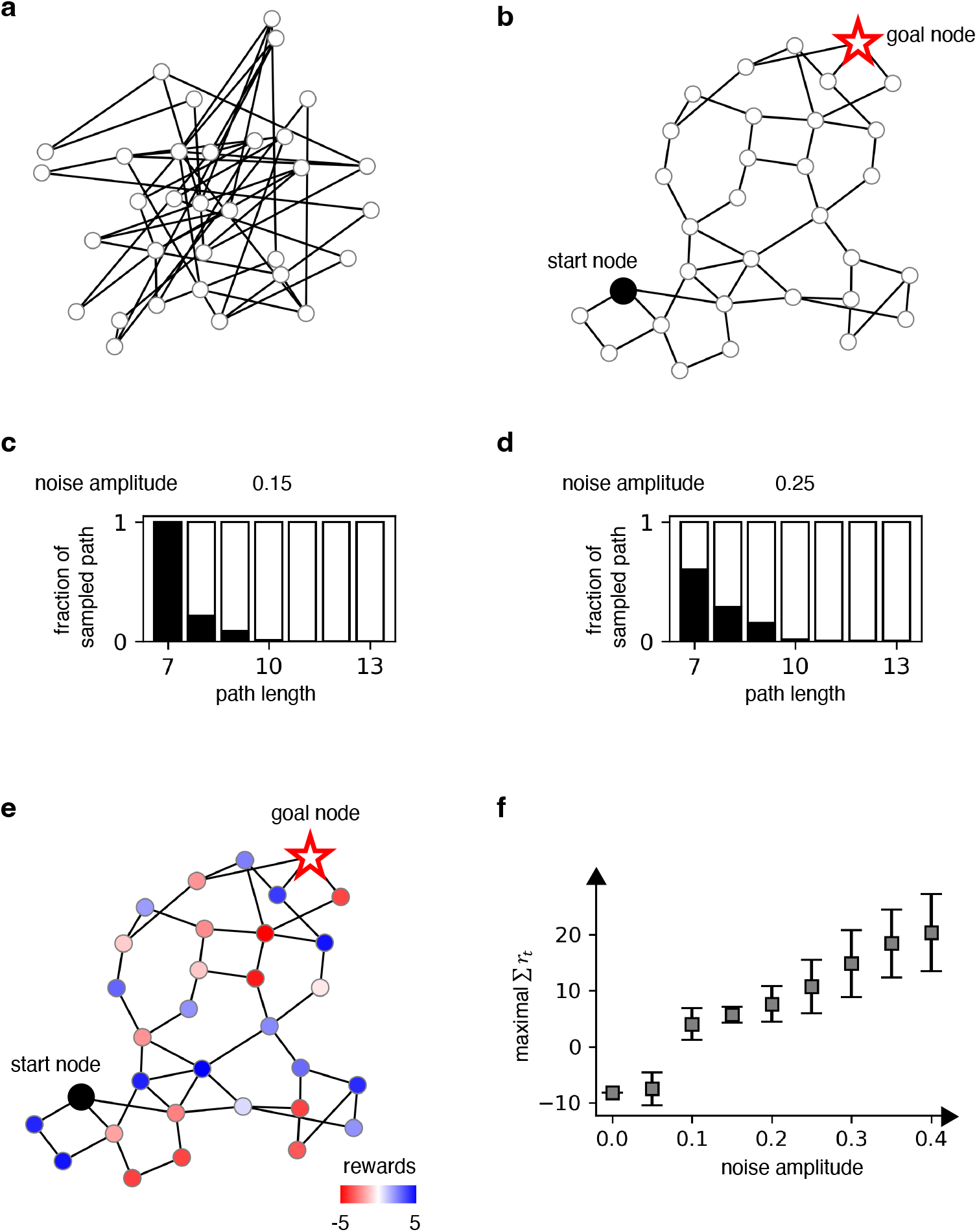
Demonstration of generic problem solving capabilities of the GCML. (a) A random graph with 32 nodes, which can be viewed as abstract model for a problem solving environment. (b) 2D projection (via t-sne) of the cognitive map that the CML creates after exploring this graph with the help of its learned embeddings of observations and actions. The geometry of this graph representation supports a simple geometric heuristic strategy for selecting actions (edges) from any given node in order to reach any distal goal node: choose an edge that points into the direction of the goal. (c) and (d) show for two levels of noise the length statistics of paths that are generated by the GCML for an arbitrarily fixed choice of start and goal node (indicated in panel b). For each path length indicated on the x-axis the black bar indicates the fraction of paths of that length from start to goal that are generated by the GCML. The lengths of paths are in both cases clustered around the minimum. The lower noise level in panel c provides a heuristic solution to the k-shortest paths problem. (e) and (f) Illustration of a functional benefit of generating a larger diversity of solution paths (with a higher level of noise in the GCML). Panel e shows a possible adhoc assignment of rewards and losses to individual nodes. If the goal is to generate a path with a maximal sum of rewards then a higher level of noise is beneficial, as shown in panel f. For each noise level 40 trajectories were generated by the GCML, and their maximal sum of rewards is indicated on the y-axis. Center points indicate mean values and the error bars indicate the standard deviation over 100 times repeated experiments with different random seeds. The starting node and goal node are the same as in b and e.

**Fig. 4:**
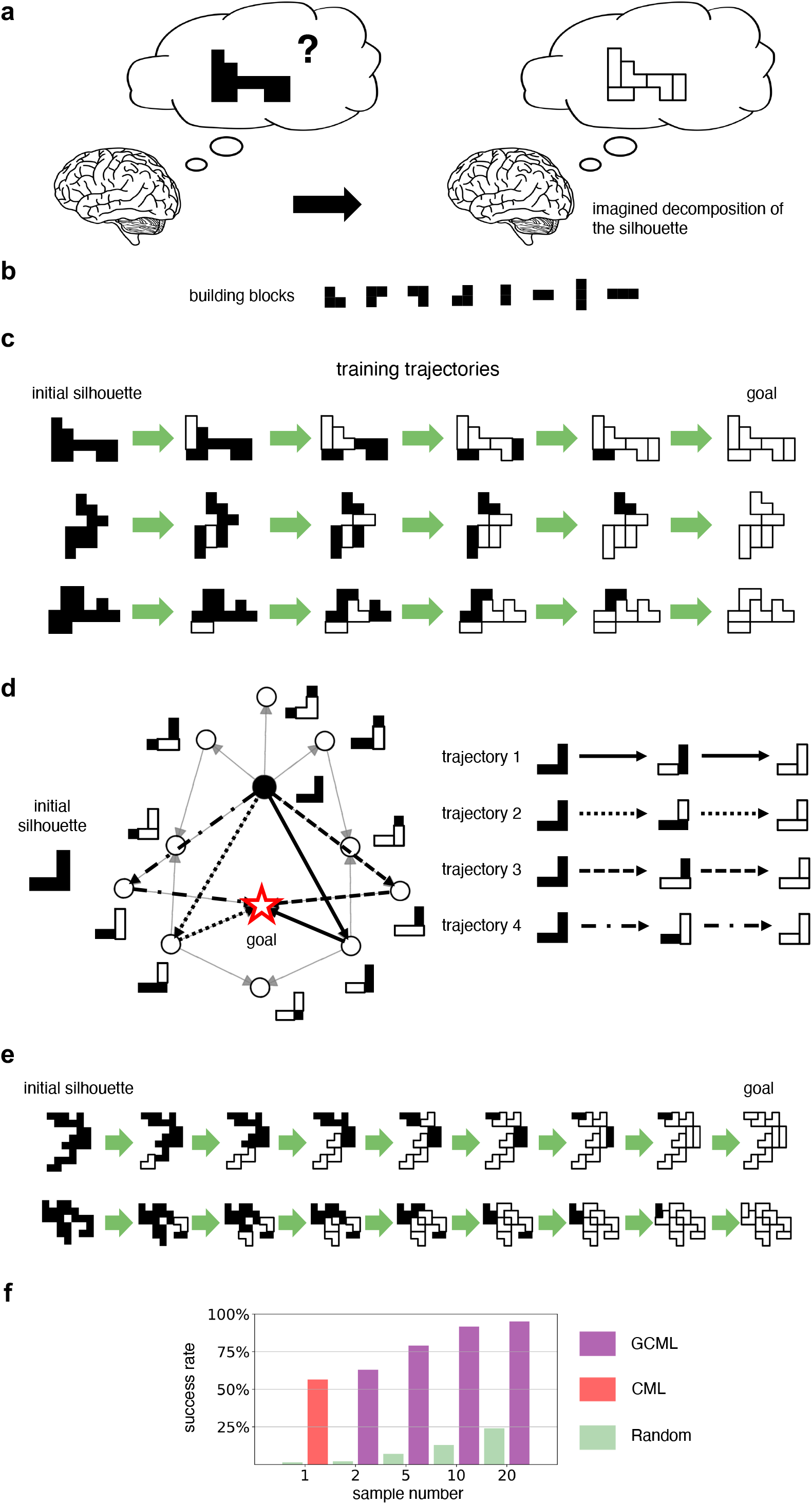
Application of the GCML to goal-directed sampling for a compositional benchmark task. (a, b) Illustration of the task: Find a decomposition of the silhouette into the building blocks shown in panel b, provided there exists such a decomposition. (c) Three example trajectories that were used during learning of the GCML. The five BBs used to generate each silhouette were removed in a randomly shuffled order. (d) 2D t-SNE projection of the resulting cognitive map for the embedding *Q* of silhouettes with affordance factors. (e) Two example trajectories generated by the GCML for decomposing silhouettes containing 8 BBs. (f) Test of the ability of the GCML to generate decompositions for decomposable silhouettes consisting of 8 building blocks, after training on decomposition of silhouettes with 5 building blocks. Hence none of the test silhouettes had been encountered by the GCML before. See Fig. S4 for additional examples.

To sum up, we have shown that stochastic sampling from a standard model for a cognitive map of 2D space supports goal-directed imagination of trajectories from start to goal locations, capturing key qualitative patterns of rodent hippocampal replay in the same setting, including generalization to un-explored parts of the space, and flexible adjustment to novel contingencies. In the following sections we will examine whether sampling from cognitive maps can also reproduce goal-directed imagination and planning in non-spatial environments.

### 2.2 Sampling from learnt cognitive maps for abstract concept spaces enables generic problem solving through goal-directed imagination

To allow navigation in (potentially) high dimensional conceptual spaces in order to solve abstract problems, see Fig. 2a for an illustration, we employ a generic model for learning cognitive maps, the Cognitive Map Learner (CML) from (Stöckl et al., 2024) that is based on the commonly accepted principle of predictive coding. As for the cognitive maps that are created through the rodent grid cell system, the key principle of the CML is to learn a forward model that relates current actions to the neural codes for the resulting states or observations. This is achieved in the CML model by learning embeddings **Q** and **V** of observations and actions into a common high-D space, see Fig. 2b. In biological terms, this high-D space models the space of neural activity patterns of neurons that encode places within the cognitive map. Hence this coding space has in the brain a dimension that is equal to the number of neurons in that area. However, a much smaller dimension – of around 1000 – suffices for the tasks considered in this study. Similarly, it suffices to employ linear maps **Q** and **V** that embed sensory inputs and efferent copies of action commands into this high-D coding space. Furthermore, simple local synaptic plasticity rules suffice for learning these embeddings in a self-supervised manner during exploration, see Fig. 2c. These local plasticity rules (see Equation 12 and Equation 13 in Section 4) approximate gradient descent for learning state predictions according to (Stöckl et al., 2024). If nonlinear embeddings would be more useful, linear readouts from fixed nonlinear circuits, as proposed by (Maass et al., 2002), could be trained equally well. A detailed description of the CML algorithm can be found in Section 4.2 and Section D of the Supplement.

Note that the CML is a deterministic model that interacts with the environment in an online manner, receiving after each action execution a resulting observation from the environment. In order to apply this model to the generation of imagined action sequences where no observations are received, we replace observations by predictions of resulting next states, see Fig. 2e. Specifically, after generating a virtual action command **a**_t_, the model does not receive any observation **o**_t+1_, and replaces it by an internally generated prediction **ŝ**_t+1_ = **s**_t_ + **Va**_t_. It continues from there to generate the next virtual action. Using this mechanism permits generating, for any given initial state **s**_t_ and any given goal state **s**\*, an imagined trajectory **s**_t_, **ŝ**_t+1_, **ŝ**_t+2_, … in the direction of the goal **s**\*, without receiving any observation. This becomes a probabilistic generative model if one adds noise in this imaginary trajectory generation process. Specifically, we have added random values from a Gaussian distribution to the current estimate of the eligibility of each action. We call this probabilistic generative model a *Generative Cognitive Map Learner (GCML)*. Note that in contrast to this generative model, the CML is a deterministic model that is not able to sample trajectories from a probability distribution. We refer to Fig. S2 and Section D of the Supplement for the detailed algorithm and a schematics of neural network implementations.

As first example for the capability of the GCML we show that it produces not just one, but a menu of possible solutions for a problem. We use here a formalization of problem solving that is widely adopted in AI and cognitive science: As task to find a short path from a start to a goal node in an abstract graph – whose nodes represent problem solving states and whose edges represent possible actions (Russell and Norvig, 2016; Newell et al., 1972; Hills et al., 2015; Geffner and Bonet, 2013). We consider a generic random graph with 32 nodes, see Fig. 3a. Whereas the CML generates according to (Stöckl et al., 2024) a single approximation to the shortest path problem, the GCML provides a selection of heuristic solutions (see Section D of the Supplement for algorithmic details). They take here the form of paths from start to goal whose length is close to minimal. Such a selection of possible solutions is useful if there are other criteria besides the length which make a particular path attractive. Mathematically, one can describe the output of the GCML as heuristic online solution to the well-known k-shortest path problem (Eppstein, 1998), where the objective is to find for some given k the k shortest paths from a given start to a given goal. As shown in Fig. 3c, most trajectories generated by the GCML in a low-noise condition (noise = 0.15) have the shortest possible length. Higher noise levels (noise = 0.25) yield a larger diversity of solutions, some of which are slightly longer paths (Fig. 3d). But their length is still clustered around the optimum. Remarkably is that they still reach the goal, in spite of the higher level of noise. Even if a bad move is initially selected, it is later compensated by moving on average in the direction of the goal. In other words, the GCML has a self-correcting homing-in heuristic to the given goal.

Computational benefits of this capability, in comparison with other algorithms fir solving the k shortest path problem, are exhibited in Fig. 6.

As illustration for a functional use of having diverse heuristic solutions to a problem we consider in Fig. 3e a case where specific rewards (blue) or losses (red) are subsequently assigned to the nodes of the graph. These could encode preferences or disadvantages of using specific nodes in a solution paths, such as avoiding on a trip a transfer at an airport where a thunderstorm is reported. We show in Fig. 3f that higher noise levels of the GCML, corresponding intuitively to more fantasy in problem solving, provide solution paths that collect a larger number of rewards, In this case we let the GCML generate for each noise level 40 trajectories, and select for each noise level the trajectory with the largest sum of rewards.

The GCML provides a neural network model for problem solving that complements previously studied ones. With the Boltzmann Machine, or via neural sampling from a spiking neural network, one had already be able to solve problems in a less flexible manner, where all constraints and desiderata are programmed into the synaptic weights of the network (Buesing et al., 2011; Habenschuss et al., 2013; Jonke et al., 2016). In contrast, the problem solving goal can be communicated to a GCML through synaptic input to the underlying neural network (see Fig. S2).

### 2.3 Sampling from cognitive maps with compositional structure enables strong generalization of problem solving

Humans are able to imagine plans to achieve truly novel goals, such as wanting to sit together with a goat on the Schafberg mountain, once they receive a verbal (or otherwise symbolic, i.e., compositional) description of such goal. In terms of cognitive maps, such goals represent states that were never encountered before during exploration. Hence the cognitive map structure around them could not be shaped by direct experience, and strong generalization from similar local motifs in the state space is needed. We wondered whether the GCML is able to reproduce this strong generalization capability of the human brain. We considered a benchmark task for problem solving in a compositional domain that was recently used in a magnetoencephalography (MEG) study of compositional planning processes in the human brain (Schwartenbeck et al., 2023). Participants were asked to decide whether a given silhouette (like the one shown in black in Fig. 4a) could be decomposed using the given set of building blocks, as shown in Fig. 4b. This compositional task domain appears to be much easier than natural language processing, but it is actually NP-hard (Moore and Robson, 2001), even for silhouettes that are just composed of the first 4 building blocks shown in Fig. 4b. This implies that there is no known determinitic algorithm that can solve this decomposition task without using computational resources that grow exponentially with the size of the silhouette.

We show here that a GCML can solve through goal-directed imagination also these quite demanding compositional tasks. A key step is that one makes the embedding **Q** of observations (here, 2D silhouettes) into the cognitive map compositional. The embedding **V** of actions (here: removal of a particular building block at a particular position of the silhouette) and the inverse model **W** are learned in a self-supervised manner from successful decomposition examples (see Fig. 4c). Decompositions that could be completed in 5 steps were used for learning. Examples are shown in Fig. 4c. The building blocks that have already been removed are indicated there in white with a black outline. This visual representation indicates the temporal decomposition process quite well, but is slightly deceptive from a mathematical perspective, because removed tiles are no longer represented in intermediate or final silhouettes. In particular, the goal state is always the same one: an empty silhouette. Hence in this case of goal-directed sampling the goal state is familiar, but the start state (the given silhouette) is in general novel.

The learnt cognitive map is for this task domain more difficult to visualize in 2D because of the large number of actions that can be applied to each silhouette (state). However, Fig. 4d shows for a simple initial silhouette that the 2D projection of the cognitive map still provides a “sense of direction” that favors actions that go into the direction of the target state (empty silhouette, indicated by a red star). Further examples are given in Fig. S4. Remarkably, after learning a cognitive map from decompositions of silhouette that are composed of 5 building blocks (Fig. 4c), the GCML is also able to decompose silhouettes consisting of 8 building blocks (Fig. 4e). Fig. 4f shows for this more demanding case, for various numbers of samples generated by the GCML, that it outperforms a baseline algorithm (“Random”) that removes building blocks randomly and the CML model. Furthermore, the GCML shows a remarkable, near perfect success rate, even with few samples.

Finally, we show in Fig. 5 that goal-directed sampling from the same cognitive map as discussed before can also solve more general types of compositional problems. One example are problems where the goal is not the empty silhouette, but a partial decomposition indicated as orange goal silhouette in Fig. 5a. Fig. 5c shows that the GCML can also solve this task, for silhouettes consisting of 6 building blocks that did not occur during learning, through goal-directed sampling from the cognitive map.

**Fig. 5:**
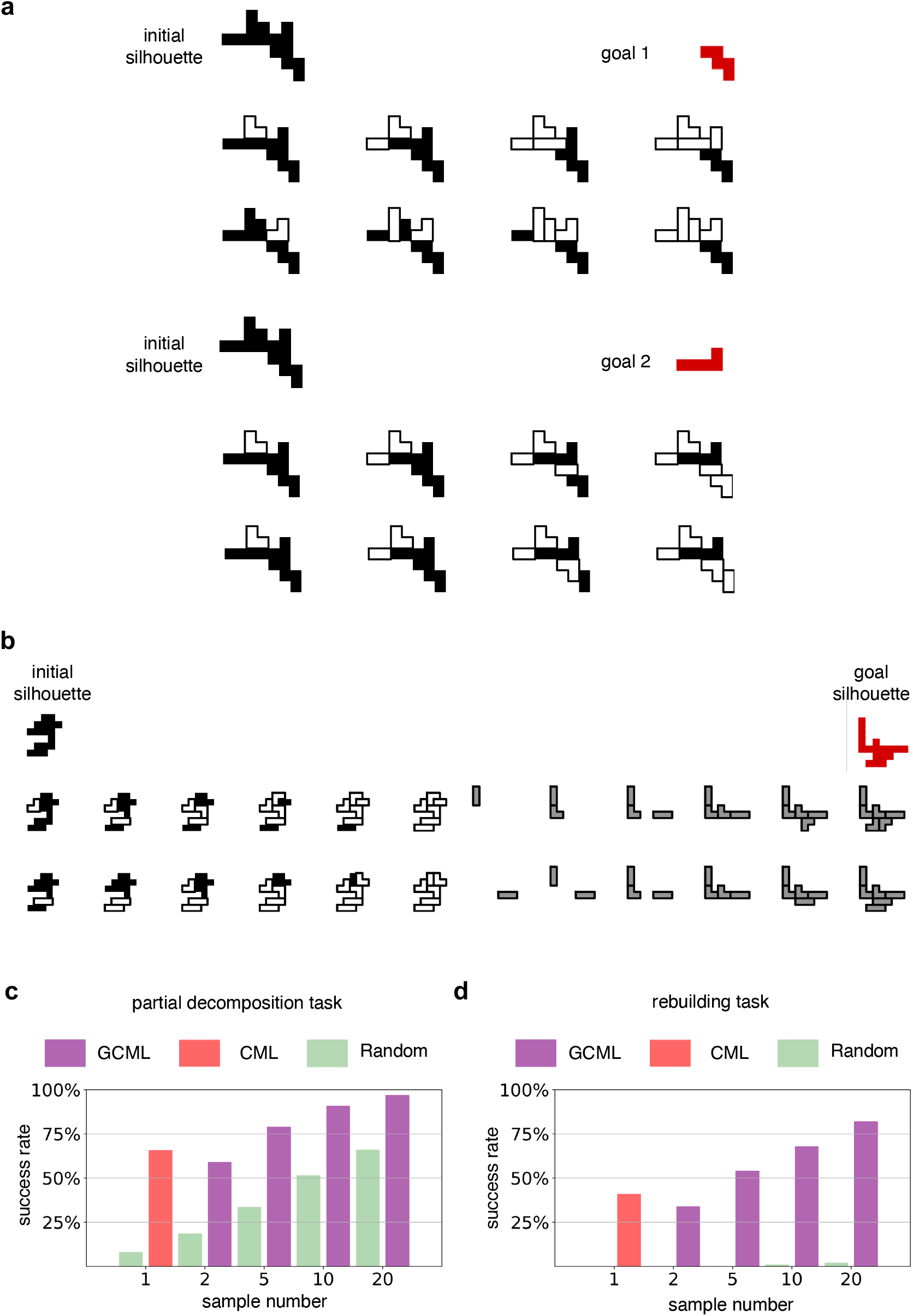
Application of the GCML for more demanding compositional computing tasks. (a) Decomposition to a specific non-empty silhouette (containing 2 building blocks, as shown in orange) from an initial silhouette that contains 6 building blocks. Successful decompositions by the GCML are shown for two examples of start- and goal-silhouettes. (b) Examples of trajectories generated by the GCML with the same cognitive map for a more demanding variant of the silhouette decomposition task: After decomposition, building blocks have to be added to form a goal silhouette. Like the initial silhouette, it contains 6 building blocks and hence did not occur during learning. Added building blocks are shown in gray for two example imagined paths to the goal generated by the GCML. (c, d) Evaluation of the performance of the GCML for both types of extensions of the silhouette decomposition task.

**Fig. 6:**
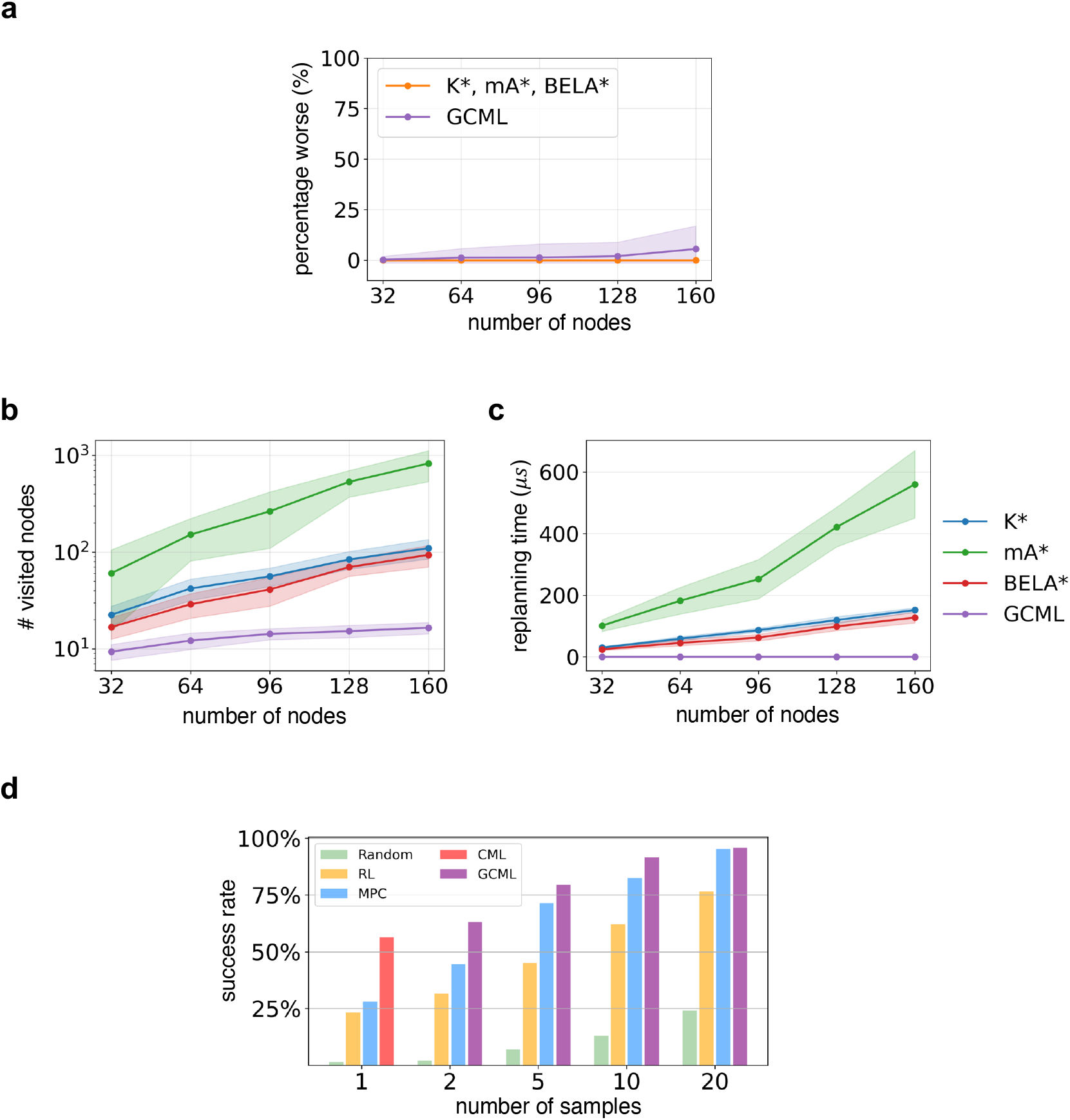
Performance comparison of the GCML with other methods for solving the k shortest path and tiling tasks. (a) **Performance for the k shortest path problem for random graphs with different numbers of nodes**. The relative percentage worse *L*_k_ of the sum of lengths of the proposed k paths is plotted for k = 5. The values for K*, mA* and BELA* coincide because they produce the optimal solution. The GCML also produces close to optimal solutions for small graphs, with a mild degradation for large graphs. (b) **Computational effort of the four algorithms**. We used the number of nodes that are visited by each algorithm to generate the first trajectory for the k shortest path problem as measure for their computational effort. One sees that the computational effort of the GCML is substantially lower than that of the other 3 algorithms. Shaded regions indicate variability across instances. (c) **Replanning latency of the four algorithms when the goal changes**. Shown is the wall clock time needed for replanning when the start or goal nodes change. The GCML incurs no replanning cost, whereas the other algorithms exhibit substantially latency caused by replanning, which increases with the size of the graph. Data in (a-c) are presented as mean ± standard deviation across 10 random graphs, with 100 random start and goal pairs per graph; the shaded error bands indicate ± standard deviation. (d) **Success rate of the GCML and 4 other methods for decomposing a silhouette into building blocks**. We consider the same task as in Fig. 4. Like in Fig. 4f we show the success rate as a function of the number of decomposition proposals that are generated. Methods include a random policy (Random), D3QN (RL), CEM-based MPC (MPC), CML, and GCML. The first RL bar (i.e., with a sample number of 1) shows the result without any noise. The performance of the GCML is substantially better when less than 20 decomposition proposals are generated.

Another example are problems where building blocks not only have to be removed, but also have to be added, in order to arrive at the goal silhouette indicated in orange in Fig. 5b. No new learning is needed for that: The action to add a building block (at a particular place of the current silhouette) is mapped to a vector that goes into the opposite direction of the one for removing this building block. Fig. 5d shows that the GCML can solve also these problems quite well, for tasks where neither the initial silhouette nor the target silhouette occurred during learning of the cognitive map.

To sum up, we have shown here that endowing the cognitive map with compositional structure permits addressing challenging problem solving tasks in compositional domains through goal-directed sampling from a cognitive map. In particular, required solution paths had to go through never encountered states. The compositional nature of the map, which reflects the relations between building blocks and their possible combinations, permits generalizing the goal-directed imagination of the GCML not only to decomposition of silhouettes that were never encountered during learning, but also to new variants of this decomposition task: partial decomposition and combining decomposition with composition of a target silhouette. Performance comparisons with other algorithmic approaches for solving tiling tasks are given in Fig. 6. Apparently the GCML is the only known online method for solving these tasks, in the sense that it can generate a first step towards a solution based on a sense of direction in the cognitive map, which could be viewed as some form of intuition, without taking the time to generate and examine a full solution path.

## 3 Discussion

We have examined a simple model that integrates three prominent facets of brain intelligence: stochasticity, cognitive maps, and compositional computing.

We have validated this model in three very different application scenarios. The first one, demonstrated in Fig. 1, reproduces biological data from the rodent brain on the imagination of possible future paths to a goal (Pfeiffer and Foster, 2013; Widloski and Foster, 2022), including rerouting around novel barriers.

Our second application has demonstrated goal-directed sampling for problem solving in abstract concept spaces. We employed for this application a generative extension of the *cognitive map learner (CML)* from (Stöckl et al., 2024), the *Generative Cognitive Map Learner (GCML)*, see Fig. 2. We have shown in Fig. 3 that the GCML provides not just a single solution to a problem, but a menu of different ones.

Finally, we have demonstrated in Fig. 4 that our model for goal-directed sampling can also be applied in compositional task domains, where a virtually infinite number of states –most of them never visited before– can be defined by new combinations of a fixed set of components. We considered there a compositional task domain that has recently been examined through recordings from the human brain in (Schwartenbeck et al., 2023), where 2D silhouettes have to be decomposed into a given set of building blocks. In Fig. 5 we have shown that the GCML can also solve demanding variations of this NP-hard compositional task, such as building a given 2D silhoutte from a given set of building blocks.

Taken together, we have shown that sampling from cognitive maps provides a surprisingly powerful and versatile heuristic method to solve computationally challenging problems across both spatial and non-spatial domains, using an algorithmic form of goal-directed imagination. Of course, there are limitations to the performance of such a heuristic online method, which generates with low latency the first step of a proposed solution. More sophisticated search, reasoning, and internal evaluation methods can be programmed into offline algorithms to achieve better performance and handle more difficult tasks. As also indicated in Supplementary Information Section A, escaping local structures can require elevated noise levels, which may reduce stability. An interesting open question is to what extent this can also be achieved with brain-inspired online methods that employs simultaneously several cognitive maps on different levels of abstraction, that also include rules and memories of preceding successful solutions to similar problems, see (Yang and Maass, 2025) for a first step.

In this work, we implemented a generative model by adding Gaussian noise to a deterministic (inverse) model, which corresponds to assuming a fixed uncertainty during sampling—a formulation commonly used in generative modeling (e.g., (Bishop et al., 1998)). Future work could explore sampling from a learned probability distribution over actions, states, and goals, which might better capture the true model uncertainty. Such an approach would connect more directly to the Bayesian framework and, more specifically, to probabilistic methods for inference and planning—such as planning as inference and active inference (Attias, 2003; Botvinick and Toussaint, 2012; Parr et al., 2022)—in which plans are inferred from learned probabilistic transition models.

Altogether we have introduced the GCML as a new model for neural sampling that supports goaldirected sampling. Hence it drastically enhances the application range of neural sampling in AI. In particular, it supports the drive towards more energy-efficient AI solutions for planning and problem solving. The local learning mechanism of the GCML makes this approach suitable for self-supervised on-chip learning during exploration. Furthermore, the (virtual) action selection mechanism of the GCML is suitable for in-memory computing with memristor crossbars, yielding a minimal delay for action selection that is independent of the size of the environment and the number of options for action selection. Hence, the GCML paves the way for creating edge devices that can employ goal-directed imagination, often referred to as fantasy, in order to find solutions to difficult and novel tasks.

## 4 Methods

### 4.1 Details of our model for spatial goal-directed imagination in Section 2.1

#### Generating a cognitive map based on grid cells

We used a simple cognitive map model, illustrated in Figure 1a, based upon the functioning of the grid cell system in the entorhinal cortex. Furthermore, in our applications, we derive place cells by combining grid firing patterns (see Section 4.1).

We have shown in Fig. 1 that these experimental data can be reproduced with a standard model for the 2D spatial map that is provided by the grid cell system of the rodent (Hafting et al., 2005; McNaughton et al., 2006; Dong and Fiete, 2024). Furthermore, an inverse model that maps differences between grid states to movement commands that reduce this difference can be learnt by a simple local rule for synaptic plasticity. By adding noise to this action selection mechanism this cognitive map becomes then a generative model that generates goal-directed movement plans that qualitatively resemble those that were decoded from the rodent brain (Pfeiffer and Foster, 2013), even rerouting around novel barriers as demonstrated in the neural recordings of (Widloski and Foster, 2022). Furthermore, the samples generated by the model generalize to never experienced trajectories, as observed in rodents (Gupta et al., 2010; Vollan et al., 2025).

The model shown in Fig. 1a consists of grid cells, place cells, and action basis neurons. All neurons were linear, with each neuron’s activation equal to the sum of its inputs. Our model is based on the core idea that grid cells can perform ideal path integration based on a velocity input (Campbell et al., 2018; Sorscher et al., 2023). We regard grid-cell activity itself as the state space of our model, in which both the target state **s**\* and its estimated current state **ŝ**_t_ are expressed, see Fig. 1b. We approximate post-training grid-cell activity in their model analytically, by a deterministic function **x** ↦ **s** ∈ ℝ ^1000^ that outputs the firing rates of 1000 grid cells at any location in a 2D arena. Each grid cell is initialized independently and modeled as the sum of three positive planar cosine gratings 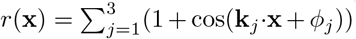, whose wave-vectors **k**_j_ are 60^°^ apart and share magnitude |**k**_j_| = 2*π/λ*. The grid scale was drawn once per cell from *λ* ~ *U* (0.05, 8) and each phase independently from *ϕ*_j_ ~ *U* (0, 2*π*). To eliminate directional bias a global rotation *θ*_0_ ~ *U* (0, 2*π*) was applied, giving arg(**k**_j_) = *θ*_0_ + *jπ/*3.

Under this setting, PCA applied to the grid-cell representations of the 10 % largest-scale (*λ*) grid cells revealed a nearly planar two-dimensional manifold (Fig. 1b). Because larger spatial periods generate progressively flatter interference patterns, this low-curvature embedding is well suited for goal-directed navigation.

Grid cell firing in the brain is updated by path integration of locomotion activity. A velocity signal is provided in our model by the activity of four action basis neurons, whose sum provides a continuous velocity vector. We constrained a fixed step size *η*_a_ = 0.05 by normalizing the length of the resulting velocity vector always to *η*_a_.

To simulate continuous movement, time was discretized, with each time step representing 10 ms. The resultant movement (Δ*x*, Δ*y*) during each time step is computed by the cumulative contributions of all four action basis neurons:

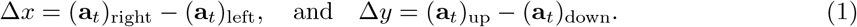

This provides the forward model for the cognitive map in this section.

As in (Sorscher et al., 2023), grid cell activity can predict place-cell activity. In our model the place fields are established by one-shot learning from grid cell activity. The incoming weight of each place cells is assigned in a single update using a simplified rule that captures the core idea of the LTP component of behavioural time-scale synaptic plasticity (BTSP) (Milstein et al., 2021; Wu and Maass, 2025).

During 500 s of random exploration, a plateau-potential gating signal occurs every 500 ms. At each event, one postsynaptic neuron *i* is randomly selected from the neuron pool of 1000 neurons, and its incoming weight vector **P**_i_ (row *i* of **P**) is overwritten with the current pre-synaptic grid-cell activity pattern **s**_t_:

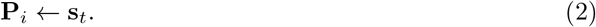

This centers the place field of neuron *i* at the location (*x*_t_, *y*_t_). Because the learning happens only at visited places, comprehensive exploration is essential; regions that are never visited cannot recruit corresponding place fields. We used grid cell activity, rather than place cell firing, for generating imagined trajectories, in particular for representing the current state of an imagined trajectory. Hence this imagination process was not affected by the lack of place cells for the unexplored region shown in Fig. 1c. Our motivation for leaving part of the 2D arena unexplored was to demonstrate that the unexplored region does not hinder grid cells from generating imagined trajectories that include segments in the unexplored region. But in order to be able to plot spiking activity of place cells during imagined trajectories in Fig. 1e we assumed that place cells had formed in the meantime also for the initially unexplored region.

#### Using the cognitive map for generating imagined paths

An inverse model in the form of a weight matrix **W** learned in a self-supervised manner to infer actions from differences in grid cell states which caused these differences, using the same simple Hebbian learning rule in Equ. 14.

We simulated here random exploration for 500 s, discretized by 10 ms within a 2D bounded environment ([0, 4] × [0, 4]), excluding the lower-left quadrant (*x <* 2, *y <* 2) shown in Fig. 1c. The agent started within the permitted region and executed a continuous-time random walk at constant speed (0.5 unit*/*s) with Gaussian turn noise for. Invalid movements (entering forbidden zones or crossing boundaries) triggered re-sampling of the direction. We consistently use the matrix **W** as learned inverse model for a cognitive map throughout this paper.

We used this inverse model for (virtual) action selection during stochastic generation of an imagined trajectories to a given goal **s**\* as follows:

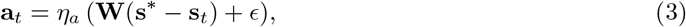

where *ϵ* is the noise that introduced stochasticity into the model, and *η*_a_ = 0.05 is the step size. Each dimension of *ϵ* was sampled from a Gaussian distribution *ϵ*_i_ ~ 𝒩 (0, *σ*_n_), where *σ*_n_ represents the “noise level” depicted in Fig. 1d. We assumed four basis action neurons defining an action vector **a**_t_ = [(**a**_t_)_up_, (**a**_t_)_down_, (**a**_t_)_left_, (**a**_t_)_right_].

Spike generation, as illustrated in Fig. 1e, was modeled by interpreting the continuous activation of each place cell *i* as a firing probability *p*_i_(*t*) within a discrete time window of 10 ms. Spikes were then sampled from a Bernoulli distribution:

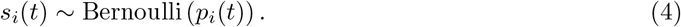

The firing probability at each time step was computed by applying the softmax function to the normalized activation *ℓ*_i_(*t*) of all place cells:

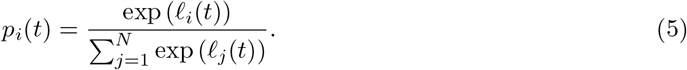

#### Obstacle avoidance in imagined trajectories

For generating imagined trajectories to a given goal in the presence of obstacles (see Fig. 1f) we augmented Equ. 3 and Equ. 1 with additional repulsive forces originating from each obstacle (*x*_o_, *y*_o_) towards the agent (*x, y*). The information encoded by the vector difference, ***δ*** = (*x*_o_ −*x, y*_o_ −*y*), may be represented, in the brain, by object vector cells (Høydal et al., 2019). Each wall was discretized into three point-obstacles (the center and two endpoints), each exerting a repulsive force **f**_i_. The total repulsive force **F** was the vector sum of these:

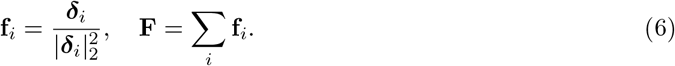

The denominator 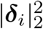 ensures the strength of each repulsive force diminishes with increasing obstacle distance. The total force **F**, scaled by factor *γ*, modified the movement of the agent during one simulation time step (10 ms), from Equ. 1 to:

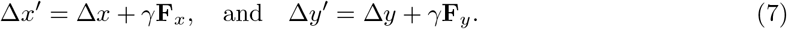

For all four navigation tasks in the presence of obstacles in Fig. 1f we set *γ* = 0.2 and noise scale *σ*_n_ = 0.3.

This provides a substantially more parsimonious model for planning 2D navigation in the presence of obstacles than the model of (Bakermans et al., 2025), where first the outer product of grid and object vector cells has to be created through an offline learning process. Our model is much simpler, and predicts that no new map has to be learnt when an obstacle is moved.

### 4.2 Details to the CML algorithm

The CML algorithm (Stöckl et al., 2024) maps observations and actions into the same high-D space. Specifically, each observation **o**_t_ is mapped using a learned embedding matrix 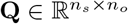 :

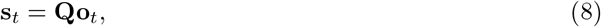

where *n*_s_ is the dimensionality of the state space, and *n*_o_ is the dimensionality of the observation **o**_t_. The target state (goal) is obtained as the embedding of a target observation:

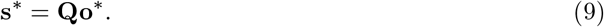

Similarly, all possible actions **a** are embedded into the same high-dimensional space via a learned embedding matrix 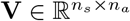, where *n*_a_ is the dimensionality of the action space. The objective of learning these embeddings is ensuring that the predicted next state **ŝ**_t+1_ is given by:

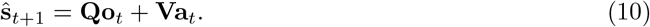

Learning aims at ensuring that **ŝ**_t+1_ becomes a good approximation of the actual next state **s**_t+1_ = **Qo**_t+1_:

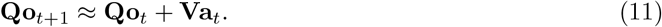

The prediction error (**s**_t+1_ −**ŝ**_t+1_), which the learning rules for **Q** and **V** aim to minimize (see Fig. 2c), has therefore the following form:

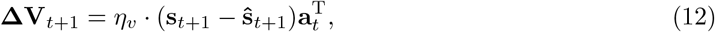

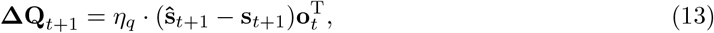

where *η*_v_ and *η*_q_ are the learning rates. These plasticity rules, known as Delta rules, approximate gradient descent to minimize the prediction error. If this learning process is successful and **ŝ**_t+1_ becomes a good approximation of the actual next state **s**_t+1_, one has a basis for generating a sequence of virtual actions in a goal directed manner, without receiving observations, as indicated in Fig. 2e.

In our implementation, the CML additionally learns during exploration an inverse model in the form of a matrix 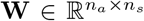. This matrix is acquired through Hebbian learning and maps state differences **s**_t+1_ − **s**_t_ onto the actions *a*_t_ that caused them (Fig. 2d):

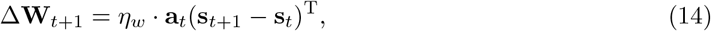

where *η*_w_ is the learning rate. **W** assigns to every possible action **a**_t_ a utility that scores its estimated usefulness (or value) for reaching **s**\* from the current state **s**_t_:

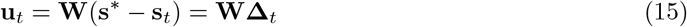

Each dimension stores the utility of a specific action, forming a utility vector **u**_t_ of length *n*_a_. The estimation of action utility is referred to as Principle 2 in CML (Stöckl et al., 2024).

This mechanism relies on the hypothesis that the geometry of the learnt cognitive map provides a sense of direction when choosing the next action. Although **W** is trained only on *local* state differences **s**_t+1_ − **s**_t_, it is applied during planning to *imaginary* state differences **s**\*− **s**_t_ that can be substantially larger. Linearity of the inverse model supports this generalization. These utility estimates are analogous to value estimates in reinforcement learning (Sutton and Barto, 2018; Sutton, 1992). In contrast to RL, however, the learned matrix **W** provides a *universal value function* that assigns values for reaching any possible state, and these estimates do not depend on an a priori chosen policy.

As in reinforcement learning, utility estimates must be modulated by an *affordance* factor that vetoes actions that cannot be executed in the current state, or that increases or decreases the utility of actions depending on their difficulty in more complex environments. The utility values are masked by the learned affordance factor 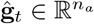, which aims to approximate the real affordance **g**_t_ defined by the constraints of the environment, a one-hot vector that indicates which actions are executable (1 for executable, 0 otherwise). This gating factor prevents the GCML from proposing infeasible actions. We denote the product of the utility of an action (for reaching **s**\*) with this affordance factor as its current *eligibility*. Action selection at time *t* then consists of choosing the action **a**_t_ with the highest eligibility.

After executing this action, the CML transitions to the next state **s**_t+1_, represented by the embedding of the next observation **o**_t+1_ (**s**_t+1_ = **Qo**_t+1_). From this new state, the same action-selection procedure is iterated until the goal state is reached.

After learning **Q, V**, and **W**, the CML can be tasked with reaching an arbitrary goal state **s**\* = **Qo**\* that results from a desired observation **o**\*. Such a target **o**\* may correspond to a spatial 15 goal location or, more generally, to a problem to solve in an abstract state space (Fig. 2a; see also (Stöckl et al., 2024)). The learned matrix **W** is then used to select actions that are likely to be useful for reaching this goal state **s**\*.

### 4.3 Mathematical description of the GCML discussed in Section 2.2, as well as its relation to biology and other theoretical models

#### Cognitive maps for non-spatial task domains

Experimental data from neuroscience and cognitive science suggest that the brain—especially the human brain—employs cognitive maps not only for spatial navigation, but also for navigation in more abstract concept spaces, where spatial locations are replaced for example by images, or by combinations of learnt ranks of an item in different linear orders (Behrens et al., 2018; Bottini and Doeller, 2020; Tavares et al., 2015; Buzsáki and Moser, 2013; Iyer et al., 2024; Xiao et al., 2025). We show here that the principles that enable according to the preceding section goal-directed imagination for 2D spatial cognitive maps generalize to cognitive maps for abstract concept spaces, and enable there generic problem solving (see Fig. 2a for an illustration of a non-spatial problem solving task).

The role of the grid-cell system in supporting the generation of cognitive maps and conceptual navigation in abstract concept spaces is now well documented across a growing body of studies, including non-invasive neuroimaging experiments in humans (fMRI/MEG; e.g., (Constantinescu et al., 2016; Park et al., 2021; Viganò et al., 2024)), fMRI studies in monkeys (Bongioanni et al., 2021), and intracranial recordings in monkeys (Veselic et al., 2025). However, the computational mechanisms that would allow grid-cell systems to scale to abstract spaces—particularly those that are not well captured by 2D Euclidean geometry and may require higher-dimensional representations—remain more debated (Dong and Fiete, 2024).

We used distinct maps to navigate physical space and abstract problem spaces. However, it is important to note that our approach to “sampling from cognitive maps” is general and can be applied across both physical and abstract spaces, provided that the cognitive map is adapted to the task and its dimensionality. Spatial navigation and foraging occur in approximately Euclidean spaces, requiring simple 2D maps. Such low-dimensional maps can also potentially be used to address simple navigation and foraging tasks in two-dimensional abstract spaces, such as the one studied in (Constantinescu et al., 2016).

However, 2D maps are insufficient for planning in non-planar random graphs and for our compositional problem solving tasks. Hence, a system like the GCML, capable of learning in high-dimensional spaces, is useful. The use of distinct maps in our tasks is therefore not motivated by the difference between physical and abstract spaces, but by the dimensionality of the problems.

Crucially, by highlighting the differences between learning maps of simpler (Euclidean, 2D) versus more complex (higher-dimensional) spaces, we are not ruling out the possibility that the grid cell system also generalizes to maps in higher dimensions required for challenging abstract tasks. For example, Dong and Fiete (2024) propose that the grid cell system can generalize to non-Euclidean environments with complex transition structures, using methods called “mixed modular coding” or “map fragmentation” (see also (George et al., 2021; Stachenfeld et al., 2017; Whittington et al., 2020) for other approaches).

By providing a general approach to sampling from cognitive maps, our method is agnostic as to whether the maps are learned through the grid cell system (an idea that is increasingly empirically supported) or with the aid of other neural systems.

#### Sampling from cognitive maps for non-spatial task domains by the GCML

From a biological perspective, the mechanism used by the GCML – the stochastic generation of multiple paths and their subsequent selection based upon expected reward – could be associated with the neural circuit formed by the hippocampus and the ventral striatum. Various studies suggest that this circuit supports goal-directed navigational planning and “vicarious trial-and-error”, with the hippocampus generating candidate future trajectories and the ventral striatum assessing the reward associated with those trajectories (Redish, 2016; Van Der Meer et al., 2012; Pezzulo et al., 2014; Pennartz et al., 2011).

The GCML uses the same cognitive maps as those that are learned in a self-supervised manner by the cognitive map learner (CML) presented in (Stöckl et al., 2024). However, the agent cannot directly access the *current* state during imagination from the observation. Therefore, the imagined state **ŝ**_t_ is used as a substitute for the actual state **s**_t_ when computing the utility:

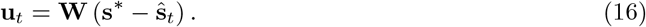

The computation of the imagined state will be detailed in the following paragraphs.

Another difference is that GCML cannot get the environment input on affordance, but the agent needs to learn and imagine it. The affordance in GCML is computed using the affordance gating matrix 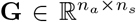 that receives the (imagined) state vector as its input. The matrix is updated during the learning process according to the following rule:

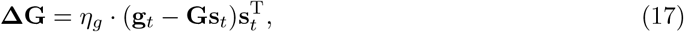

where *η*_g_ represents the learning rate.

The resulting vector, after applying the affordance gating, is termed the eligibility:

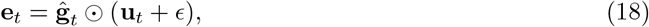

where *ϵ* is the noise is a crucial term in GCML that turns the CML into a probabilistic generative model. This injected noise enables GCML to explore actions beyond the one with maximal eligibility. Hence, in contrast to CML’s deterministic selection, GCML stochastically generates diverse trajectories.

Finally, the selected action **a**_t_ is the one with the highest eligibility:

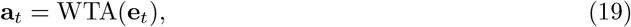

where WTA(·) denotes the winner-take-all operator, selecting the action with the highest eligibility from all possible actions. It can be approximated by a simple neural circuit with lateral inhibition (Nessler et al., 2013).

After selecting a virtual action **a**_t_ at the estimated state **ŝ**_t_, the estimated next state, *t* + 1, is calculated using the bootstrapping process:

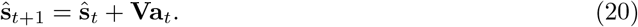

By iteratively applying the trajectory generation process described earlier, a virtual goal-directed trajecvtory is generated.

By repeatedly generating trajectories in the presence of inherent noise in Equ. 18, different trajectories are generated. This results in a set of possible solutions from the initial state **s**_0_ to the goal **s**\*.

A downstream network could select a specific one from this repertoire of imagined trajectories based on a given criterion. For example, when rewards are associated with specific states, the estimated rewards 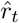 of each state can be accumulated over an entire imagined trajectory, and the one with the largest sum of predicted rewards can be selected (see Fig. 3f).

#### The GCML suggests a new model for neural sampling

On a more abstract algorithmic level the GCML provides an intriguing novel paradigm for using the inherent noise of biological and physical computing systems as a computational resource (Maass, 2014). A new feature is that this noise enables goal-directed sampling, for a goal that is provided in the form of a synaptic input, rather than just sampling from a fixed probability distribution that has been programmed into the weights of synaptic connections. This endows the neural sampling method with a brain-like flexibility in adjusting to new goals or contingencies, and to solve new problems that were never encountered before. From a more general theoretical perspective our model for goaldirected sampling is related to previous models for probabilistic inference in the brain for perception and action selection through sampling (Pouget et al., 2003; Fiser et al., 2010; Ujfalussy and Orbán, 2022; Parr et al., 2022; Ma et al., 2006; Friston, 2010). But in contrast to this preceding work it provides a neural network paradigm for sampling from a marginal distribution, relative to problem specifications that are defined through the current synaptic input to the network. A theoretical basis for that had been provided in (Habenschuss et al., 2013). It was shown there that a stationary distribution exists for this sampling from a marginal distribution even if the synaptic connections of the network are not symmetric, which is always the case if one has synaptic connections between excitatory and inhibitory neurons. It also exists if the neurons are spiking, and exhibit a brain-like diversity of firing properties.

From a biological perspective, our approach is supported by experimental evidence showing a dual role of (hippocampal) cognitive maps, for learning states in an associative mode and supporting sequential activity in a predictive mode (Liu et al., 2023) – and provides a possible mechanistic explanation of this finding. novel empirical predictions that can be addressed in future studies. In particular, our model proposes that the brain learns together with each cognitive map also an inverse model that maps state differences to possible actions for reducing this state difference. Note, however, that an inverse model for a cognitive map could also be implemented in the brain in a somewhat different way, where the current and goal states are provided separately as inputs, instead of their difference. But as long as the inverse model is approximately linear, it supports local actions with “foresight”, preferring actions that move into the direction of the goal.

#### Predictions of the model tests through biological experiments

Future studies could align this proposal with empirical findings showing that goal and reward information shape trajectory representations and the geometry of cognitive maps in the hippocampus and cortex (Gauthier and Tank, 2018; Muhle-Karbe et al., 2023; Hok et al., 2007; Schuck et al., 2016; Crivelli-Decker et al., 2023; Nyberg et al., 2022; Zutshi et al., 2025; Bernardi et al., 2020). The assumption that goal-directed behavior is driven by minimizing the discrepancy between current and goal states is shared with frameworks such as cybernetics (Rosenblueth et al., 1943; Wiener, 1961; Powers, 1973) and active inference (Parr et al., 2022).

A key prediction of the GCML model for future biological experiments is that noise in the selection of possible actions during imagination is directly related to the resulting diversity of action plans, and implicitly also on the difficulty of solving a given task. This could be tested experimentally.

Furthermore, our model predicts that the speed of the generation of the next step of a multi-step solution plan depends only mildly on the current distance to the goal. The reason is that the next step is generated by a feedback circuit as indicated in Fig. 2e whose processing speed is independent from the distance to the goal. But the generation speed of the next step is predicted to grow with the number of action alternatives that are available, since this is likely to affect the speed of the underlying WTA computation for each action selection.

Finally, our model predicts that current goals can be decoded from brain areas that are involved in action selection, as already partially shown in (Zutshi et al., 2025). In contrast, RL models predict that the values of adjacent states are most relevant for action selection, and the identity of the current goal has no direct influence.

#### GCMLs and RL

Sampling from a marginal distribution provides an alternative to RL for generating goal-directed behavior. Models based on RL often require methods from machine learning for training, such as BP and BPTT, whose biological viability is debated. Furthermore, except for methods based on successor representations, RL approaches require retraining for each new goal that is to be reached. The approach of (Stachenfeld et al., 2017) used the successor representation for RL in combination with a cognitive map. We are not aware of efforts to sample from this data structure. It is also an open problem whether this approach can be made applicable to tasks such as the compositional tasks that we consider, where planning needs to visit states that were not encountered during exploration.

This also holds for Single-Goal-Conditioned-Contrastive RL (SGCRL), that was recently proposed (Liu et al., 2024; Bastankhah et al., 2025). But this approaches focuses, like the CML and GCML, on learning state representations that support fast action selection. In fact, actions are selected there also through a WTA operation applied to a dot product of vectors, one of which represents the goal. The focus of SGCRL is on producing efficient exploration strategies, however for a fixed goal. An open question is whether its method could also be used to enhance goal-invariant exploration in cognitive map based approaches.

Apart from the versatility advantage of cognitive map based methods, there are also some recent experimental data which suggest that dopamine signals support learning of various types of predictions in the brain (Jeong et al., 2022; Gershman et al., 2024; Kahnt and Schoenbaum, 2025), not only reward predictions. Hence they might also support learning of cognitive maps by minimizing prediction errors for sensory inputs, which provides the basis for the CML and GCML model. Altogether, the diversity of data on dopamine signals suggests that the brain could employ not just one but a multitude of mechanisms and strategies for learning goal-directed behavior, as suggested by (Gershman et al., 2024).

#### GCMLs and other theoretical models

There are various models for how the brain could learn cognitive maps, with a particular emphasis on the hippocampal–entorhinal system. Some approaches, such as the TEM (Whittington et al., 2020) and others (Chandra et al., 2025), emphasize a separation between learning the general structure of the environment (putatively via a grid code)—for example, the transitions between spatial locations, which generalize across different mazes—and learning the content of specific locations (putatively via place cells), which is maze-specific. Other models dispense with this distinction (George et al., 2021; Raju et al., 2024) but are still able to learn, offline, the transition structure of complex, non-Euclidean spaces.

The GCML, in contrast, can learn cognitive maps in high-dimensional spaces through a simple online, predictive learning rule. A unique feature of the GCML (and its predecessor, the CML) is that the learned embeddings provide a sense of direction toward goals—a capability not explicitly addressed in the aforementioned methods or related approaches.

Model Predictive Control (MPC) is another already existing framework for action selection based on a learned dynamics model. Compared with the GCML it has the advantage that it can handle also continuous state and action spaces. But apparently it lacks the capability of the GCML to choose goal-directed actions with short latency, that result from the embedding of states and actions into a high-D space. A nice challenge for future research will be to combine the best of both approaches in a hybrid model.

### 4.4 Further details for the application of the GCML for goal-directed imagination for generic problem solving (Section **2.2**)

The task setup was a random graph with 32 nodes, where the degree of each node was uniformly sampled between 2 and 5. Observations **o**_t_ corresponded to nodes, and actions **a**_t_ represented traversing edges in a specific direction. Hence each edge is modeled by two possible actions. Both **o**_t_ and **a**_t_ were encoded as unique one-hot vectors. The training dataset consisted of 200 randomly generated trajectories, each of length 32. During the planning process, GCML samples a set of trajectories from a given starting node to a target node.

The mappings **G** transform the estimated high-dimensional state **s**_t_ into corresponding affordance factors. The affordance gating vector is a binary vector indicating available actions, where ones represent feasible actions and zeros indicate restricted actions. A linear transformation is sufficient in this setting, as the vectors **s**_t_ are approximately orthonormal.

Exploration was regulated by adding a Gaussian noise vector ***ϵ***—with each dimension sampled from 𝒩 (0, *α*_ϵ_)—to a normalized unit-length utility vector. The noise amplitude *α*_ϵ_ was set to a default value of 0.1.

Initial values for model parameters were drawn from Gaussian distributions: **Q** ~ 𝒩 (0, 1) and **V, W, G** ~ 𝒩 (0, 0.1). Smaller initial values for **V** improved performance. Learning rates were set as *η*_q_ = 0.1 and *η*_v_ = *η*_w_ = *η*_g_ = 0.01. GCML demonstrates robustness to variations in learning rates, similar to the original CML, with stable training performance for *η*_g_ in the range [0.001, 0.05].

Each node was assigned a reward drawn from a uniform distribution 𝒰 (− 5, 5).

To demonstrate that GCML provides a good approximation of the top-k shortest path problem (see Figs. reffig:fig3c-d), 40 trajectories were generated from a given starting node to a goal node under different noise levels in the GCML. The number of distinct trajectories for each possible trajectory length (i.e., the number of edges in the path from the starting node to the goal node) was computed using a breadth-first search algorithm. For each trajectory length, the proportion of distinct trajectories generated by a given noise scale GCML relative to the total number of possible distinct trajectories serves as a measure of how well GCML approximates the top-k shortest path algorithm.

### 4.5 Details of goal-directed sampling in compositional task domains, discussed in Section 2.3

This domain is arguably more transparent than natural language, but it still exhibits a core feature of compositional computing, its computational complexity. In fact, this decomposition task is NP-hard, which implies that every known deterministic algorithm requires computational resources that grow exponentially with the problem size(Moore and Robson, 2001; Russell and Norvig, 2016). Recent experimental data suggest that the human brain solves it by addressing possible building blocks in a sequential manner (Schwartenbeck et al., 2023). Their recordings from the human brain showed that during planning, participants considered the possible building blocks that could be removed from the silhouette in a sequential manner, starting with the most obvious candidates. Furthermore, the authors suggested that replay sequences supported a hypothesis testing process for silhouette decomposition.

Other experimental studies demonstrate more generally that replay from spatial and non-spatial cognitive maps provides a neural substrate for goal-directed imagination and compositional computing. But few studies have so far advanced mechanistic explanations of these phenomena (Foster, 2017; Stoianov et al., 2022; Bakermans et al., 2025; Kurth-Nelson et al., 2023; Wittkuhn et al., 2021). The silhouette decomposition problem involves finding a set of building blocks (BBs, or “tiles”) that completely cover a given silhouette without gaps or overlaps. This process is called the decomposition of the given silhouette. GCML generates candidate decompositions through goal-directed sampling, similar to the previously described navigation tasks. In the simplest version of the problem, the goal node **o**\* is defined as the empty silhouette, while the starting node **o**_0_ is the silhouette to be decomposed. An action is defined as removing a BB from a particular position in the remaining silhouette. Details of this process are provided below.

#### Building Blocks

We used the same set of building blocks as considered in the experiments of (Schwartenbeck et al., 2023), except that we removed one of the building blocks (an atomic square tile) that would make every silhouette decomposable. Hence there were eight predefined building blocks (BBs). The first four BBs were categorized as *corners*, each with four possible orientations and a width of 1 pixel. The remaining four BBs were *bars*, including two longer types (1× 3 and 3 ×1 pixels) and two shorter types (1× 2 and 2 ×1 pixels). These eight BBs were derived from the human silhouette decomposition experiment of (Schwartenbeck et al., 2023). But we excluded the 1 ×1 BB since each presence makes each given silhouette decomposable. GCML was tasked with imagining possible decompositions of given silhouettes, each composed exactly of five BBs.

Observation of a silhouette is encoded by a binary vector that has a 1 at each pixel position in a 2D grid space that corresponds is black in the silhouette.

The simplest possible compositional embedding **Q** of silhouettes is the identity map over these binary vectors. This provides already decent performance of the GCML. However, its performance increases substantially, and its cognitive map is endowed with a better sense of direction if the embedding **Q** slightly departs from the identity map and assigns greater weight to black pixels that are surrounded in a silhouette by white pixels. This embedding increases the utility of removing building blocks that cover black pixels which protrude from a silhouette. This heuristic is beneficial since the choice of these building blocks is more constrained and therefore less error-prone. It also creates a cognitive map of the compositional domain that supports the same geometric heuristic for action selection as in the previously discussed GCML applications: Choose an edge from the current node that best points into the direction of the goal, see Fig. 4d. Further details and the full algorithm are provided in Methods and part C of the Supplement.

#### Generation of Training and Test Datasets

Observation of a silhouette was encoded by a binary vector that has a 1 at each pixel position in a 2D grid space that corresponds is black in the silhouette. Each silhouette in the dataset was created by randomly selecting five BBs and placing them within a 10 ×10 image at random positions. In total, 18,000 training samples and 2,000 test samples were generated. Any silhouette appearing in the training dataset was excluded from the test dataset, regardless of its position. Each sample represented a decomposition trajectory, in which BBs were sequentially removed from an initial silhouette. The silhouette generation process followed these rules: all BBs had to remain within image boundaries without overlapping. An initial BB was placed randomly within the image with equal probability for each position. Subsequent BBs were added adjacent to the existing silhouette, with at least one pixel adjacency to increase task difficulty. Longer adjacencies increased the number of possible removal actions, further challenging the model. In the training dataset, the five BBs composing each silhouette were removed sequentially in random order to generate training trajectories.

#### Details of Actions

In the silhouette decomposition problem, each action **a**_t_ corresponds to removing a specific BB from a specific valid position within the 10 x 10 grid, resulting in 664 possible actions. Each action is uniquely encoded as a one-hot vector. This formulation aligns with human experiments (Schwartenbeck et al., 2023), where participants manipulate BBs using a mouse and keyboard to cover the silhouette. Since moving the same BB to different positions requires distinct movements, each BB at a different position is treated as a separate action.

#### Details of Observations

In the silhouette decomposition problem, the observation **o**_t_ ∈ ℝ^10×10^ directly represents the silhouette image, rather than being encoded as a one-hot vector as in previous experiments.

#### High-Dimensional Embedding Details

The simplest possible compositional embedding **Q** of silhouettes is the identity map over these binary vectors. This provides already decent performance of the GCML. However, its performance increases substantially, and its cognitive map is endowed with a better sense of direction if the embedding **Q** slightly departs from the identity map and assigns greater weight to black pixels that are surrounded in a silhouette by white pixels. This embedding increases the utility of removing building blocks that cover black pixels which protrude from a silhouette. This heuristic is beneficial since the choice of these building blocks is more constrained and therefore less error-prone. It also creates a cognitive map of the compositional domain that supports the same geometric heuristic for action selection as in the previously discussed GCML applications: Choose an edge from the current node that best points into the direction of the goal, see Fig. 4d. The state representation **s**_t_ = **Qo**_t_ lies in a 100-dimensional space that matches the dimensionality of the observation. In our implementation, however, this effect is achieved entirely through the contextdependent affordance term (see the paragraph describing g_2_), which already assigns higher influence to protruding pixels through its convolution-based computation. As a result, no explicit modification of **Q** is required, and we maintain **Q** = **I**. Further algorithmic details are provided in Supplementary Section C.

#### Details of learning the cognitive map

The action embedding matrix **V** is learned according to Equ. 12. The learning rate of **V** was set to *η*_v_ = 0.01, with values between 0.0001 and 0.1 also demonstrating stable performance. In this silhouette decomposition task, each action corresponds to removing a specific building block at a specific valid location, and the corresponding action embedding **V** is trained to predict the resulting silhouette after the removal of that block (Equ. 11).

The inverse model **W** is learned in a self-supervised manner from successful decomposition trajectories using Equation 14 with a learning rate *η*_w_ = 1, and the weights in **W** saturate at 1:

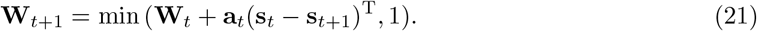

As a result, **W** becomes a binary matrix. Both matrices **V** and **W** were initialized using Gaussian distributions 𝒩 (0, 0.001). The method remained effective across a broad range of initial standard deviations, from 0.0001 to 1.

One can easily see that **W** outputs for a given state difference for each possible action (BB) the number of pixels in the overlap between the state difference and the BB, provided that the BB is contained in the state difference (otherwise it outputs 0).

#### Affordance Details

The affordance gating mechanism in the silhouette decomposition problem comprises two components. The first component, g_1_, ensures the agent selects only BBs present in the silhouette. We assumed GCML could directly access g_1_ from the environment during all imagined steps. This assumption aligns with biological experiments (Schwartenbeck et al., 2023), where subjects directly observe the screen while solving the silhouette decomposition problem. Thus, affordance information is assumed available at all steps, requiring no inference from **ŝ**_t_.

The second component, g_2_, represents context-dependent affordance determined by the estimated state **ŝ**_t_. Each pixel, affordance value is set as the number of non-empty adjacent pixels (top, bottom, left, and right). This is modeled as a convolution operation with the kernel:

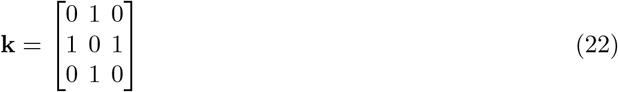

using a 3 × 3 kernel size, padding of 1 with value 1, and stride of 1. The context-dependent affordance function g_2_ transforms the delta vector **Δ** = **s**\* − **ŝ**_t_ into:

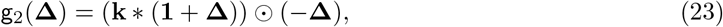

where ∗ denotes the convolution, and ⊙ represents the element-wise multiplication. The delta vector **Δ** contains unremoved pixels with a value of −1 and all other pixels with a value of 0. The expression (**1** + **Δ**) flips these values, converting −1 to 0 and 0 to 1, so the subsequent convolution counts the number of adjacent pixels with a value of 0. Finally, since only the affordance values at unsolved pixels are relevant, an element-wise multiplication between the minus delta vector (−**Δ**) and the affordance value **k** · (**1** + **Δ**) is performed as the second component of the affordance check.

#### Trying to decompose a silhouette

This task was formulated as a navigation problem, where the embedding of the empty silhouette, **o**\* = **0**, represented the target node in the high-dimensional state space (cognitive map), while the embedding of the initial silhouette, **o**_0_, served as the starting node. In the presence of noise, GCML selected the action with the highest utility value from the set of currently feasible actions, determined by the environment’s affordance factors g_1_(·) and g_2_(·). This selection process followed the directional intuition of the cognitive map:

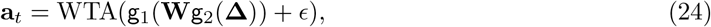

where the noise term *ϵ* followed a Gaussian distribution 𝒩 (0, 0.1).

After selecting an action **a**_t_, the next state was estimated as **ŝ**_t+1_ = **ŝ**_t_+**Va**_t_, and GCML proceeded to the next iteration to attempt the removal another BB. The algorithm terminated when no feasible actions remained. If all pixels were removed, GCML successfully identified a valid decomposition; otherwise, the attempt was deemed unsuccessful.

The four paths from start to goal that are plotted in Fig. 4d with solid, dotted, dashed and dashdotted line segments correspond to four trajectories that solve the problem. Their intermediate states are indicated at the right of Fig. 4c. Note that all of these correct trajectories result from actions that move into the direction of the goal in this cognitive map. The top part of the left plot in the panel depicts three silhouettes that result from removing a “wrong” building block from the starting silhouette that does not lead to a full decomposition of the initial silhouette. Note that they lie relative to the starting point on the opposite direction of the goal. Another dead end configuration, that arises if one does not move towards the goal node in the second removal step of the paths indicated by solid and dotted lines, is shown at the bottom. Additional examples of cognitive maps and generated trajectories to decompose silhouettes are shown in Figs. S4.

#### Details of performance evaluation

The performance of GCML was evaluated based on the proportion of test cases in which it successfully solved the given silhouette in the test dataset.

As baseline comparisons, we consider a random strategy that removes as many BBs as possible from the silhouette in a random manner, as well as the CML algorithm (Stöckl et al., 2024).

#### Decomposition to a given non-empty goal silhouette

In the non-empty goal silhouette decomposition problem, GCML removes BBs from a given silhouette **o**_0_ to transform it into various non-empty target silhouettes **o**\* ≠ **0**. When generating the test dataset, the initial silhouettes contained six BBs, generated following the same method described earlier. Any initial silhouette that appeared in the training dataset was removed, regardless of its position. The goal silhouettes contained two BBs, generated by randomly removing four BBs from the six BBs used to generate the initial silhouettes. GCML could plan trajectories for non-empty goal silhouettes using Equ. 24 without requiring retraining. The performance was evaluated using 1,000 test silhouette pairs.

#### Combining sihouette decomposition and composition

To address the silhouette rebuilding problem, GCML began with a given silhouette and iteratively modified it by adding or removing BBs to reconstruct the goal silhouette. This task was solved using the same cognitive map learned from the silhouette decomposition problem introduced earlier, leveraging mathematical symmetry.

GCML followed a two-stage approach. In the first stage, it removed BBs as long as possible from the given silhouette using a straightforward method. Here, GCML adopted for that a sub-goal **s**^†^ = **0**, and the delta vector was computed as **Δ** = **s**^†^ −**s**_t_. Then it switched to the second stage, where it aimed at constructing the given target silhouette by sequentially adding BBs. More precisely, the goal was the final silhouette **s**\* = **Qo**\*, and the delta vector was given by **Δ** = **s**\* −**s**_t_. Since adding BBs was the inverse operation of removing them, GCML selected actions based on:

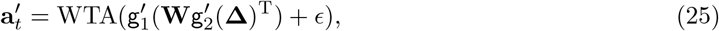

where the affordance factor 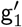 ensured that GCML only added BBs contained within the goal **s**\*. The function 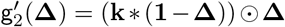 was designed following the same principle as in Equ. 23. The negative transformation of **Δ**’s sign in Equ. 23 facilitated the addition of BBs as the inverse of their removal. Here, 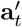 represented the action of adding a BB corresponding to **a**_t_, which removed a BB.

For the testing dataset, the initial and goal silhouettes were generated using the five BBs following the previously introduced generation process. To test the robustness of our model, a randomly selected noise-pixel was added to the same empty location in both the start and goal silhouettes. Any initial or goal silhouette that appears in the training dataset was removed, regardless of its position. A total of 1,000 initial and goal silhouette pairs were generated to test the performance.

### 4.6 Comparison of the performance of the GCML with competing methods for solving the k shortest path and tiling tasks

We compared the performance of the GCML on the k shortest path problem with that of three other algorithms that have been proposed for this task: The K*, mA*, and BELA* algorithms. The results are shown in Fig. 6 a-c. In Fig. 6 d we compare the performance of the GCML for tiling tasks with that of a RL algorithm and the MPC algorithm. We found that the performance of the GCML is quite competitive, in spite of the fact that it is an online algorithm: It immediately generates a first heuristic step towards a solution, without generating a complete solution path, or even generating and comparing many possible solutions paths in an offline manner, as the competing methods do (we are not aware of competing online methods for solving this task, except for the Random method examined in Fig. 6 d). Our comparisons show that the GCML requires altogether much less computational effort and latency. We also expect that it is superior when fast adaptation to a variation of the task is desired.

Detailed description of the competing algorithms and details of our comparisons are given below. We also discuss there differences in the amount of working memory that is required by different approaches.

#### 4.6.1 k shortest path problem

As shown in the main text, GCML is an approximate k shortest path algorithm. We compare it with three k shortest path algorithm baselines: the classical K* (Aljazzar and Leue, 2011), mA* (Flerova et al., 2016), and a more advance bidirectional edge labeling A* algorithm (BELA*) (Linares López and Herman, 2024). None heuristic form of the algorithms is used, since nodes in our random graphs environment are equivalent.

##### Details of K* algorithm

K* is an on-the-fly algorithm for enumerating the k shortest paths in nondecreasing cost order (Aljazzar and Leue, 2011). It first runs an A*-style search to obtain the shortest path and to incrementally reveal sidetrack edges (i.e., detours) relative to the current shortest path tree. These detours are organized into a path graph on which a Dijkstra-like procedure is executed to repeatedly extract the next best detour combination, reconstruct the corresponding full path, and output it. The two searches are interleaved: when the auxiliary search exhausts currently known detours, K* resumes A* to expand more of the original graph and updates the auxiliary structure, continuing until k solutions are produced.

##### Details of mA* algorithm

mA* (Flerova et al., 2016) extends A* (Hart et al., 1968) from returning a single shortest path to enumerating the k shortest paths in nondecreasing path length. It maintains a priority queue of candidate nodes initialized with the start node and repeatedly selects the most promising node according to an evaluation that combines the accumulated path length with a heuristic estimate of the remaining distance to the goal (i.e., the remaining distance is fixed at zero when no heuristic information is provided). The selected node is expanded to generate successors (i.e., child nodes). The algorithm discards any successor that has already been generated k times; otherwise, the successor is inserted into the priority queue with its updated cost. Unlike standard A*, it does not terminate after the first goal-reaching path; each time a complete start-to-goal path is found, it is output as one solution and the search continues until k solutions are produced.

##### Details of BELA* algorithm

Compared with mA*, BELA* (Linares López and Herman, 2024) does not enumerate the k shortest path by repeatedly pushing many alternative path-specific copies of the same state into the priority queue. Instead, BELA* keeps the forward search closer to a single A*-like exploration that builds an explored graph with sufficient predecessor information, and it delegates top k enumeration to a separate layer: it stores each newly certified deviation from the shortest-path tree as a centroid (i.e., a compact representation of a family of solutions) and later reconstructs multiple complete paths at once by combining compatible prefixes and suffixes from the explored structure. Consequently, BELA* emphasizes implicit representation of many solutions and outputs them only when their ordering among the next shortest solutions is guaranteed.

#### Measure of performance

To quantify the performance *L*_k_ of a k shortest path algorithm, we compute the relative error in the total length of the returned k paths. Let *p* denote the summed path length of the different k shortest paths produced by the algorithm, and let *p*_k_ denote the ground-truth summed path length. We define *L*_k_ as

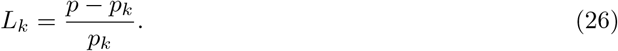

Lower *L*_k_ is better.

We also report the latency to obtain the first shortest path to the goal, measured as the number of visited nodes required to generate the first trajectory. For the baselines, this metric counts how many times a node is removed from the candidate queue and expanded by generating successors and inserting them into the queue. It is a standard measure of search effort and an implementation-robust proxy for computational cost. Baseline methods typically visit far more nodes than the length of the returned trajectory because they must explore the graph. For GCML, each decision selects an affordance-feasible action and transitions to the next state, so the visited-node count equals the sampled trajectory length.

#### Comparison of performance

Here, we set *k* = 5 for all experiments. We evaluate all methods on 10 graphs for each node size (*N* ∈ {32, 64, 96, 128, 160}) and 100 random start and goal pairs per graph. The graph structure and the start and goal pairs keep unchanged for all algorithms. For GCML, to ensure a well-trained cognitive map at larger *N*, we fix the number of trajectories to 1,000 and the state dimensionality to 5,000; all other settings follow the Methods section.

Results are shown in Fig. 6a,b. As shown in Fig. 6a, the three baselines are exact and therefore achieve *L*_k_ = 0. GCML degrades slightly as *N* increases. However, as shown in Fig. 6b, GCML visits far fewer nodes to generate the first potential trajectory of the solution than the baselines, benefiting from the directional guidance provided by the cognitive map, which allows it to avoid extensive exploration of the graph as baseline algorithms.

#### Comparison of replanning latency

We also measured replanning latency when the start or goal node changes while keeping the graph fixed. Replanning latency is defined as the wall-clock time an algorithm needs to explore the graph after the start or goal query is changed while the underlying graph remains fixed. All experiments related to wall-clock time in this section were run on a workstation equipped with dual Intel Xeon E5-2690 v4 CPUs (2.60 GHz), providing 28 physical CPU cores and 56 hardware threads in total, and 6 NVIDIA Quadro P6000 GPUs connected via PCIe, with 24 GB of on-board memory per GPU. As shown in Fig. 6c, the replanning time of the baseline algorithms increases with the number of nodes. In contrast, because GCML relies on the learned cognitive map, it incurs negligible additional replanning cost.

#### Comparison of working memory consumption

We define the working memory of the algorithms as the transient memory required during online trajectory generation. The three search-based baselines impose a substantially heavier online memory burden than GCML, albeit for different reasons. K* must preserve structures for sidetrack management and path enumeration. mA* must maintain a dynamically growing search frontier, a record of explored states, and multiple predecessor references to support the recovery of alternative shortest paths. BELA* also requires a record of labeled explored state and auxiliary structures called centroids for combining prefixes and suffixes of candidate paths. As the number of requested solutions increases, these intermediate representations of the baseline algorithms can accumulate.

By contrast, GCML performs online generation by following a learned cognitive map prior and sampling directly toward the goal, so its transient memory is dominated by the current local decision context and the partial trajectory being produced, rather than by a global search over the graph. Consequently, GCML’s online working memory is typically much closer to a local, near-constant-size inference buffer, whereas the baselines require graph-scale, search-dependent memory that grows with problem difficulty and the number of requested paths. In conclusion, GCML may incur a fixed static memory cost for storing model parameters and the learned cognitive map, whereas the working memory of the search baselines grows dynamically during online computation.

#### 4.6.2 Tiling problem

We implemented two mature and widely used baselines on the same tiling environment setting used by our method (i.e., five BBs in training silhouettes and eight BBs in the test silhouettes): a reinforcement learning (RL) baseline, and a model predictive control (MPC) baseline.

#### Details of RL algorithm

For RL algorithm, the dueling double deep Q-Network learning (D3QN) algorithm is adopted. The Q-function uses a dueling architecture (Wang et al., 2016) with Double-DQN target selection (Van Hasselt et al., 2016). The definition of observation and action keeps unchanged to GCML. When selecting actions, the action feasibility is enforced by an affordance mask as in GCML (invalid actions are masked before selection by affordance). We tuned the learning rate over {10^−3^, 10^−4^, 10^−5^, 10^−6^} and selected the value with the best validation performance. We also varied the depth of the encoder’s linear layers, the value and advantage heads, adopting the best-performing architecture.

The network model consists of a convolutional encoder, and two dueling heads for value and advantage estimation. The encoder has (i) a convolutional stack with channels 1 →32→ 64 →64, using 3 × 3 kernels with stride 1 and padding 1, followed by ReLU activations, and (ii) a feedforward stack 6400 →1024 →1024 →512 with LayerNorm and GeLU activations, producing a 512-dimensional state representation *s*. On top of this representation, the value head is an multilayer perceptron (MLP) with layer size 512 →512→ 256→ 128 →1 that outputs *V* (*s*), and the advantage head is an MLP 512 →512 →256→ 128 →|𝒜| that outputs *A*(*s, a*); both heads use LayerNorm and GeLU activations. 𝒜 denotes the set of all possible actions.

We train the network with a dueling critic and experience replay. The Q-network is optimized with Adam (learning rate 10^−5^) using mini-batches of 512 transitions sampled from a replay buffer of capacity 2 × 10^5^. We use a discount factor *γ* = 0.99. Training is run for 10,000 episodes. Exploration follows an *ϵ*-greedy policy with *ϵ* linearly annealed from 0.30 to 0.02 over 1,200 episodes. During evaluation, to generate different trajectories, we keep a stochasticity with *ϵ*_eval_ = 0.3 (optimized among {0.05, 0.1, 0.15, …, 0.4}). With probability *ϵ*_eval_, it performs policy-guided stochastic exploration: affordance-masked action scores from the trained Q-network are converted into a probability distribution via softmax, and a valid action is sampled from this distribution. The per-step reward is designed to reward the valid decisions: *r*_t_ = 2 · 𝕀_done_, where 𝕀_done_ indicates task completion.

#### Details of MPC algorithm

To choose an action, MPC (Garcia et al., 1989) plans an optimal action sequence under a learned policy model, executes only the first action, and replans at the next step by discarding the remaining actions. In our implementation, MPC uses the cross-entropy method (CEM) (Kobilarov, 2012) as its policy-optimization (planning) procedure. CEM maintains a sampling distribution over action sequences and updates it iteratively over time. At each iteration, it draws candidate sequences, rolls them out through the learned dynamics to approximate the induced state trajectories and returns, and scores each candidate accordingly. A fixed number of the best-scoring sequences are then treated as elites, and their statistics are used to refit the distribution, yielding an updated action distribution for the current decision step.

Further details of the MPC algorithm are given below. The observation and action definition is same as GCML. At real step *t*, MPC optimizes an action sequence

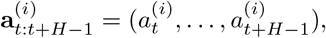

for *i* = 1, …, *N* sampled rollouts, where *H* is planning horizon and *N* = 96 is sample count. We set the horizon *H* to 5 to stay consistent with training on 5-BB silhouettes, and then test performance on 8-BB silhouettes.

Sampling is performed from per-time-step categorical distributions:

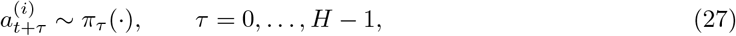

with affordance like GCML. Here, *π*_τ_ (·) is categorical policy at planning step *τ*. After simulating a candidate sequence, we compute the CEM score defined as

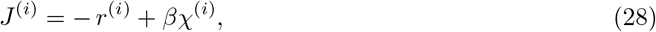

where 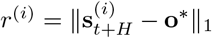 is the terminal mismatch; *χ*^(i)^ = 𝕀 [*r*^(i)^ = 0] is the success flag; 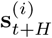is the remaining silhouette at step *t* + *H*; **o**\* is goal silhouette; *β* = 100 is terminal success bonus. Higher *J*^(i)^ is better.

For each planning step *τ*, the elite empirical distribution is

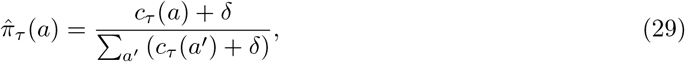

where *c*_τ_ (*a*) is elite count of action *a* at step *τ*, and *δ* = 10^−6^ is the minimum action probability. Then distribution smoothing is

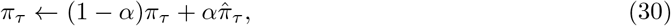

followed by clamping *π*_τ_ (*a*) ≥ *δ* and re-normalization. Here *α* = 0.3 is smoothing factor.

CEM optimization runs for *I* = 4 iterations, then only the first action of the best sequence is executed:

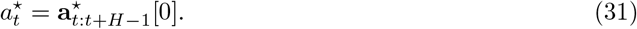

One BB is removed following 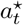, horizon shifts by one step, and planning repeats until success or no more BBs could be removed.

#### Comparison of performance

The success rates (defined in the same way as for GCML) for the random baseline, RL, MPC, CML, and GCML are shown in Fig. 6d. All algorithms are evaluated on the same 2,000 test samples as described in the Methods section.

For the RL baseline, the network architecture is much more complex than those of the other methods. It also requires backpropagation to train, which makes deployment on neuromorphic hardware challenging. In addition, the reward is difficult to design, and the sparsity of the reward signal makes training unstable.

For the MPC baseline, there is no policy training, and it achieves higher success rate than the RL baseline. However, compared to RL and GCML, MPC incurs higher planning cost because it performs an iterative sampling-and-refinement procedure at every decision step, rather than amortizing computation across episodes. In our implementation, MPC uses CEM to optimize a horizon *H* action sequence by repeatedly rolling out *N* candidate sequences for *I* iterations, which results in a per-step workload proportional to *O*(*I*· *N* ·*H*). This issue is exacerbated in compositional tiling tasks where early placements can create irreversible geometric constraints, so short-horizon optimization may fail to anticipate long-term feasibility. Consequently, when the sampling budget is reduced (smaller *N* or fewer iterations *I*), MPC’s distribution updates become noisy and can collapse to suboptimal modes, leading to a pronounced drop in success rate. In contrast, GCML leverages a cognitive map that provides a global embedding of the observations and the learnt utility, which guides sampling toward regions that are more likely to yield feasible completions. Therefore, under comparable or even smaller online budgets, GCML can maintain higher success and better sample-efficiency than MPC in current setting. Overall, we emphasize that MPC remains a strong general-purpose control baseline, but its per-step compute and horizon-limited look a head make it less attractive for the highly combinatorial environments like tiling problem. Same as RL baseline, it also needs to define a rollout score Equ. 28 to measure the quality of a candidate action sequence.

Our GCML achieves the best overall success in this task across different sample numbers and requires algorithmically simpler local updates compared with RL and MPC.

## Data availability

Our work does not utilize any specific dataset, making it independent of proprietary data sources.

## Code Availability

Code used for training and evaluating GCML is publicly available via GitHub at https://github.com/LH-cbicr/GCML and Zenodo at https://doi.org/10.5281/zenodo.19370442 (ref.Lin and Yang (2026)).

## Acknowledgments

HL and RZ were supported by the Brain Science and Brain-like Intelligence Technology-National Science and Technology Major Project 2021ZD0200300. YY and WM were partially supported by the National Science Foundation of the USA (EFRI BRAID project 2318152) and the Austrian Science Fund (FWF) (10.55776/COE12). GP was partially supported by funding from the European Research Council under the Grant Agreement No. 820213 (ThinkAhead), the Italian National Recovery and Resilience Plan (NRRP), M4C2, funded by the European Union – NextGenerationEU (Project IR0000011, CUP B51E22000150006, “EBRAINS-Italy”; Project PE0000013, “FAIR”; Project PE0000006, “MNESYS”), and the Ministry of University and Research, PRIN PNRR P20224FESY and PRIN 20229Z7M8N. This research was funded in whole or in part by the Austrian Science Fund (FWF) 10.55776/COE12. For open access purposes, the author has applied a CC BY public copyright license to any author accepted manuscript version arising from this submission.”

## Author contributions

HL, YY, and WM designed the models and the tasks. HL and YY carried out numerical simulations, and analyzed the results. HL, YY, RZ, GP, and WM wrote the paper

## Supplementary Information

### A Obstacle avoidance of goal-directed sampling from a cognitive map based on grid cells (complementing Section 2.1 and Fig. 1)

**Fig. S1:**
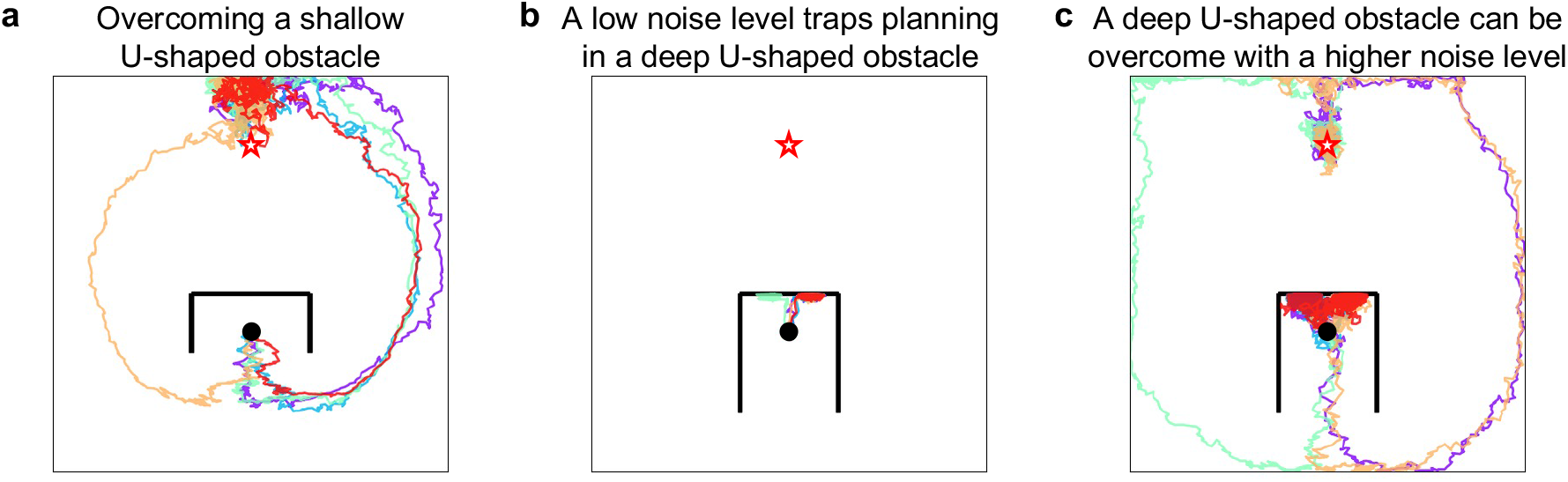
A sufficiently large noise level enables planning to escape from U-shaped obstacles. (a) With the same noise level 0.3 as in Fig. 1f the goal-directed sampling can overcome shallow U-shaped obstacles. (b) This does not hold for a deeper U-shaped obstacle. (c) But with noise level 0.5 the same goal-directed sampling can overcome also the deeper U-shaped obstacles, and still reach the goal.

In this section, we evaluate the ability of goal directed sampling from a grid-cell based cognitive map to handle navigation tasks that require detours. As example we consider escape from U-shaped obstacles.

As described in Section 4.1, the agent’s movement results from the combined effect of three components: (1) attraction toward the goal, (2) stochasticity introduced through the noise term *ϵ* in action selection, and (3) a repulsive force exerted by obstacles. The sum of these components determines the direction of each movement step.

Detour-based navigation, where the agent must temporarily move away from the goal, is enabled by the latter two components: noise and repulsion. Using the same parameters as in Fig. 1f, we generated the trajectories shown in Fig. S1a. When the U-shaped obstacle is shallow, the repulsive force alone is sufficient to drive the agent around the obstacle; all paths consistently move downward from the starting point.

However, this simple mechanism fails when the obstacle becomes deeper (Fig. S1b). In this case, the extended lower walls exert upward repulsive forces, which—under the simple repulsion rule used in Section 4.1, trap the agent within the U-shaped obstacle.

Increasing the noise level alleviates this issue (Fig. S1c). Higher stochasticity allows the agent to occasionally escape the U-shaped region; once outside, the repulsive force becomes beneficial again, pushing the agent away from the obstacle and toward the goal. Thus the noise level regulates to what extent goal-directed planning can overcome obstacles by transiently moving away from the goal.

This simple planning method provides an alternative to the navigation strategy of (Bakermans et al., 2025). Note that even more difficult obstacles can be handled with higher dimensional cognitive maps as considered in Section 2.2. But there are certainly also limitations to the problem solving capability of a simple heuristic online algorithm that is equipped with just a single cognitive map. For example, to escape from an U-shaped obstacle one would need higher and higher levels of noise when the U-shape becomes deeper. Then a hierarchical organization becomes more appropriate, where a problem is divided on the top level into subproblems that each require different solution methods. An interesting open problem is whether a more generally approach with several hierarchically structured cognitive maps could achieve that.

### B Possible implementation of the generation of cognitive maps by CMLs and GCMLs

The cognitive map is generated for the GCML in the same way as for the CML. The latter was described in detail in (Stöckl et al., 2024), and two network diagrams are repeated here for the convenience of the reader.

The architecture consists of several populations of neurons. Peripheral neurons encode observations (blue in Fig. S2), and other neurons encode actions, using one-hot coding (red in Fig. S2).

Delay modules introduce one-time-step delays through inhibitory relays. Such delay modules are readily available in neuromorphic hardware platforms, including Spinnaker and Intel’s Loihi chip (Davies et al., 2021), as well as for in-memory computing systems.

As illustrated in Fig. S2, the model learns the parameters **Q, V** through minimizing the prediction error **Qo**_t+1_ − **Qo**_T_ − **Va**_t_ (Fig. S2(a)), while **W** is updated via Hebbian plasticity (Fig. S2(b)).

**Fig. S2:**
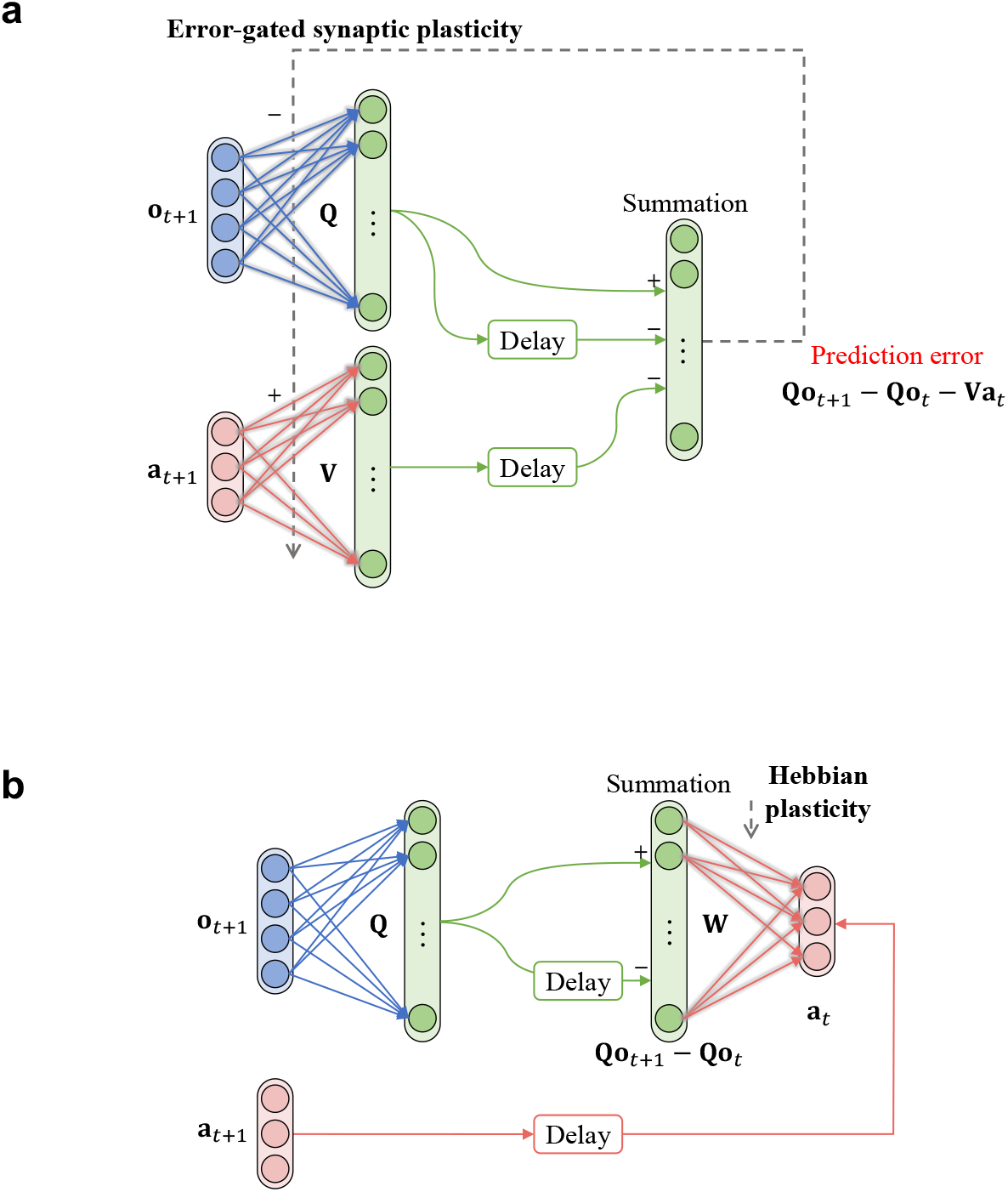
Generation of cognitive maps for the CML and GCML through self-supervised learning (copied from Stöckl et al. (2024)). **a** Network structure for cognitive map learning. The prediction error, computed by the population of linear units on the right, gates synaptic plasticity between neuron populations representing observations, actions, and internal states of the cognitive map. **b** Network structure for learning the weight matrix **W** (inverse model) using Hebbian plasticity. Some signals pass through inhibitory interneurons (not shown), indicated by a negative sign at the corresponding synaptic connections. These signals are assumed to be delayed by one time step.

### C Scaling properties of the CML and GCML

We scale up the graph size and average path length, and also the dimension of the state space, in order to see how the performance of the GCML degrades.

We first examine how the performance of the CML and the GCML degrades when the average shortest-path length increases. The results are shown in Fig. S3a as functions of the number of nodes in the graph. The average shortest-path length between node pairs increases from 2.90 to 7.60 (green curve and y-axis on the right). Performance is quantified as the relative path-length increase over the Dijkstra shortest path:

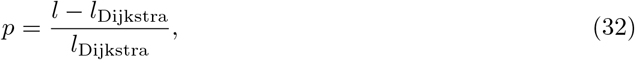

where *l* is the path length generated by the CML or GCML, and *l*_Dijkstra_ is the shortest-path length computed by the Dijkstra algorithm. For each graph size, 100 random start–goal pairs are evaluated. Note that the performance of the GCML degrades significantly more slowly than that of the CML. The dimension of the high-dimensional state space is set to 3,000 for both models. Exploration is carried out using 2,000 trajectories, each consisting of 128 steps. For the GCML, the number of sampled trajectories is fixed to 40 and the noise scale is fixed to 0.15. All other settings match Section 4.4. To generate random graphs with *N* nodes, in contrast to Section 4.4, we first constructed a ring structure by connecting nodes sequentially, and then randomly added *N/*4 additional edges between randomly selected node pairs. This process allows to control the sparsity of the graph, while making sure that there exists a path between any pair of nodes. A sparser graph leads to longer average shortest path length between all pairs of nodes in the graph.

#### Impact of the dimension of the state space on performance

We fixed the number of nodes at 128 and varied the dimensionality of the high-dimensional state space. As shown in Fig. S3b, GCML performance improves as the state dimension increases from 25 to 3,000. All other parameters remain identical to those in the path-length scaling experiment. Performance and standard deviations are computed over 100 random start–goal node pairs.

#### Scaling of GCML performance in approximating the *k* shortest-path problem

As described in Section 2.2, the GCML provides an approximate solution to the *k* shortest-path problem. Here, we evaluate how the quality of these approximate solutions degrades as the number of nodes in the graph increases. The results are shown in Fig. S3c,d. In both the 96-node graphs (Fig. S3c) and the 256-node graphs (Fig. S3d) the dimension of the state space of the GCML is fixed at 3,000, and all other parameters remain unchanged. According to Fig. S3c, the GCML performs well for four randomly selected node pairs in 96-node graphs, with shortest-path lengths ranging from 7 to 14. However, when the number of nodes increases to 256 (Fig. S3d), GCML fails to generate any valid shortest path of length 10 or 18 between the given start and goal nodes when the noise scale is 0.15. Increasing the noise scale to 0.25 alleviates this issue to some extent, but the GCML still does not generate all possible short path lengths.

### D Details of the GCML planning algorithm for sections 2.2 and 2.3

Algorithm 1 provides all details of the GCML algorithm for the abstract graph task, and Algorithm 2 for the silhouette decomposition task. All notations are consistent with those used in the Methods section.

**Fig. S3:**
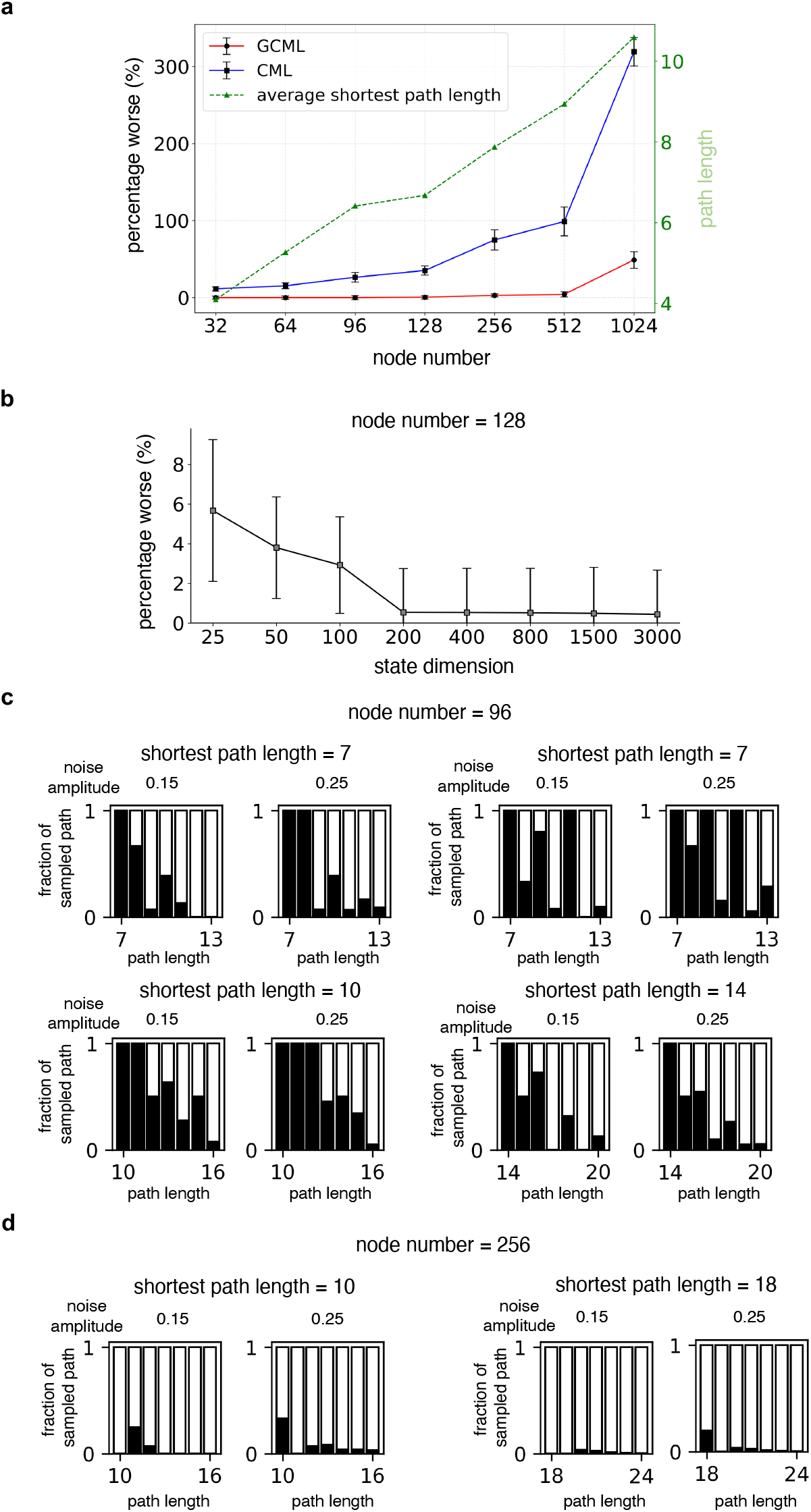
Scaling of the GCML performance on random graphs. (a) Increase of the minimal length of paths produced by the CML and GCML with increasing size of the random graph for random start and goal nodes, in comparison with the shortest path produced by the Dijkstra algorithm. One sees that the performance of the GCML degrades substantially more gracefully. (b) Dependence of the minimal length of paths (again in comparison with the shortest path produced by the Dijkstra algorithm) that are produced by the GCML for a random graph with 128 nodes, shown here as function of the dimension of the state space. One sees that performance improves when the dimension of the state space increases. Mean percentage worse is shown by the center points, and standard deviations are shown by the error bands, computed over 100 random start–goal node pairs in (a) and (b). (c, d) Distribution of the lengths of paths produced by the GCML for random graphs with increasing number of nodes. As in Fig. 3 c,d the length of the black bar indicates for each path length the fraction of paths of that length from the given start to the given goal that are generated by the GCML. Panel c shows results for graphs with 96 nodes, and panel d for graphs with 256 nodes. In comparison with Fig. 3c, d for random graphs with 32 nodes one sees here that a decent approximate solution to the k-shortest path problem is still provided for graphs with 96 nodes, but not for graphs with 256 nodes.

#### Algorithm 1

Generating Trajectories in the Abstract Graph Task

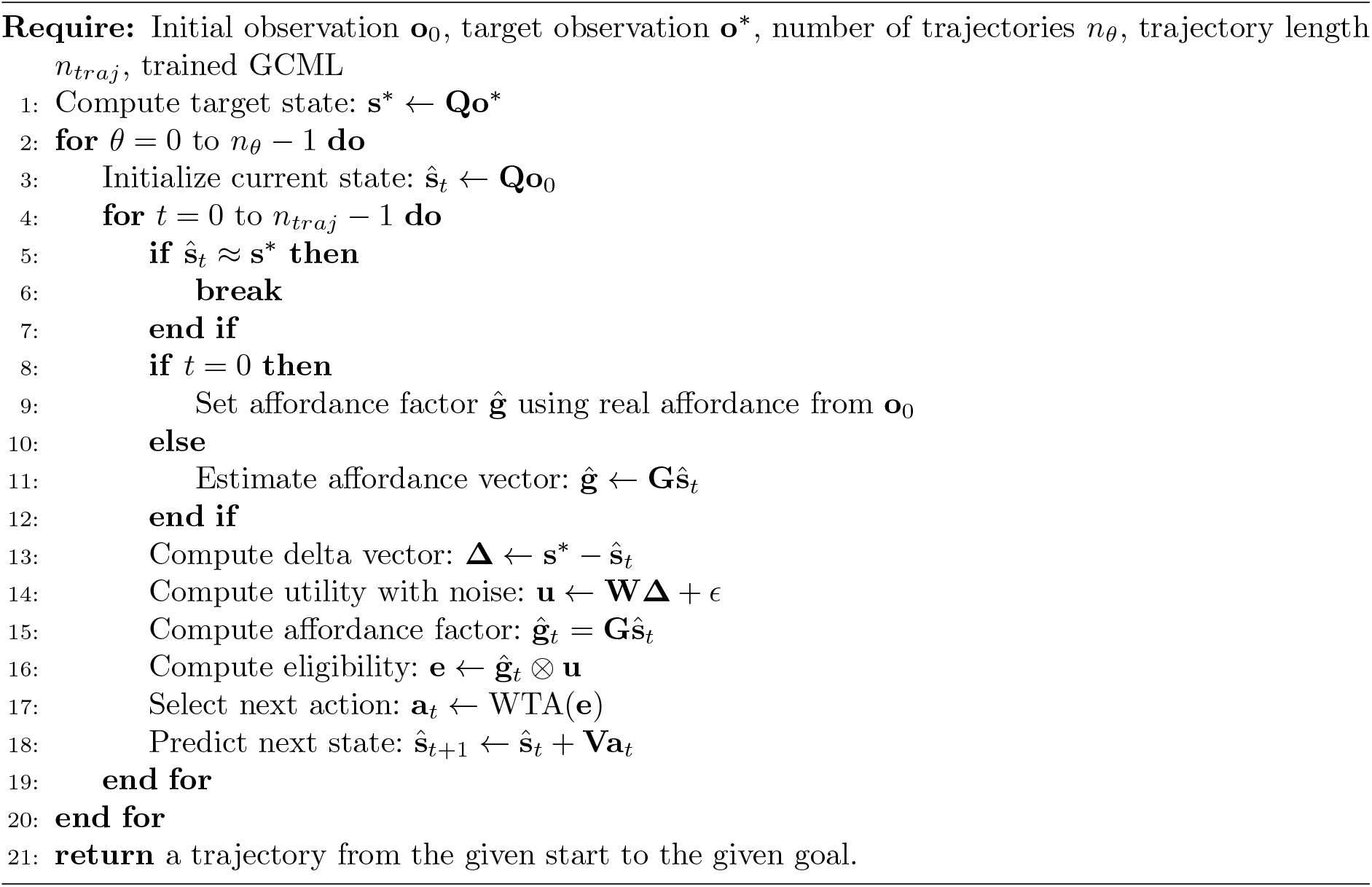

#### Algorithm 2

Generating a Set of Possible Decompositions in the Silhouette Task

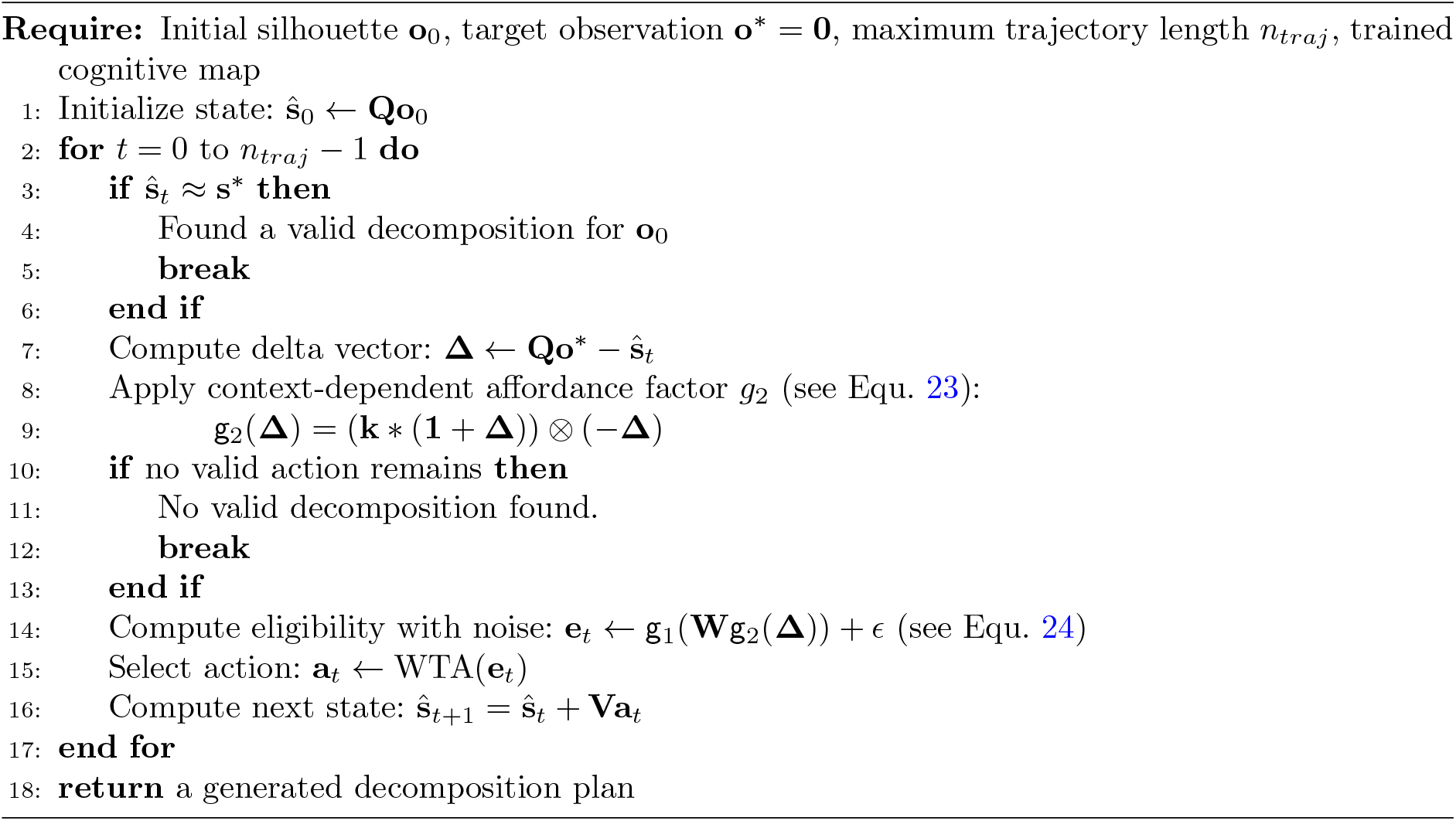

### E Cognitive maps for two further instances of the tiling-decomposition task considered in Section 2.3

**Fig. S4:**
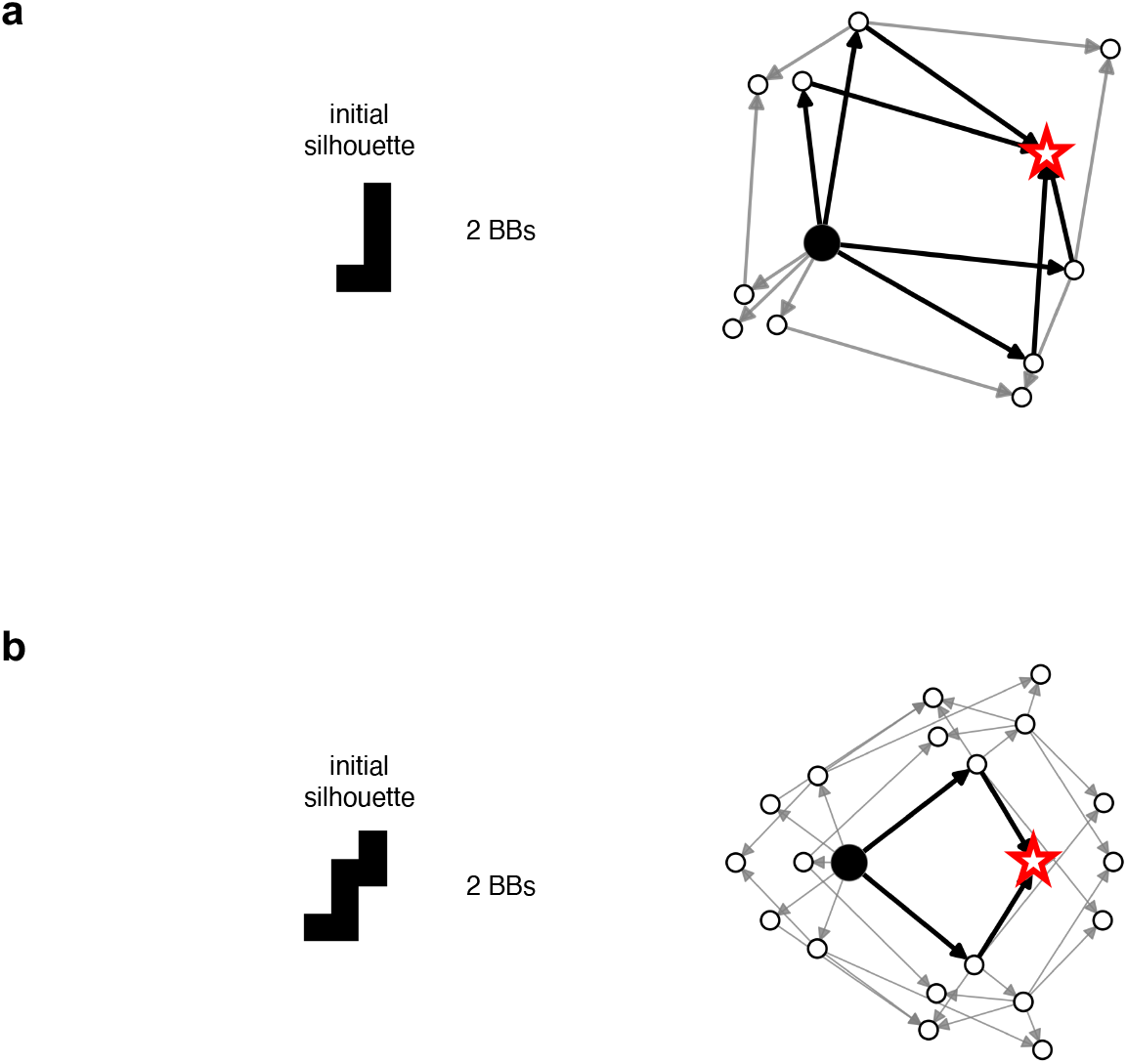
Two further examples for the cognitive map that the GCML generates for the tiling problem. Panels (a) and (b) show the cognitive map of a silhouette containing two BBs. The left panels display the initial silhouette, and the right panels present 2D projections of their corresponding cognitive maps. The black node represents the high-dimensional state **s**_0_ of the initial silhouette, and the red star denotes the high-dimensional state **s**\* of the empty goal silhouette. Each arrow corresponds to the embedding of an action (i.e., removing a BB from a silhouette). All nodes are calculated following Eq. 23. All 2-D projections are generated via t-SNE (PCA initialized). The solid black arrows indicate trajectories that successfully decompose the initial silhouettes.

Fig.S4 shows 2D t-SNE projections for two further instances of the compositional task addressed in Fig.4. One sees that also for these examples the cognitive maps provide a sense-of-direction for decomposing the given silhouette.

## References

Aljazzar, H. and S. Leue. 2011. K∗: A heuristic search algorithm for finding the k shortest paths. Artificial Intelligence 175 (18): 2129–2154.

Attias, H. 2003. Planning by probabilistic inference. In International workshop on artificial intelligence and statistics, pp. 9–16. PMLR.

Bakermans, J.J., J. Warren, J.C. Whittington, and T.E. Behrens. 2025. Constructing future behavior in the hippocampal formation through composition and replay. Nature Neuroscience: 1–12.

Bastankhah, M., G. Liu, D. Arumugam, T.L. Griffiths, and B. Eysenbach. 2025. Demystifying the mechanisms behind emergent exploration in goal-conditioned rl. arXiv preprint 2510.14129.

Behrens, T.E., T.H. Muller, J.C. Whittington, S. Mark, A.B. Baram, K.L. Stachenfeld, and Z. Kurth-Nelson. 2018. What is a cognitive map? organizing knowledge for flexible behavior. Neuron 100 (2): 490–509.

Bernardi, S., M.K. Benna, M. Rigotti, J. Munuera, S. Fusi, and C.D. Salzman. 2020. The geometry of abstraction in the hippocampus and prefrontal cortex. Cell 183 (4): 954–967.

Bishop, C.M. 2006. Pattern recognition and machine learning. Springer.

Bishop, C.M., M. Svensén, and C.K. Williams. 1998. Gtm: The generative topographic mapping. Neural computation 10 (1): 215–234.

Bongioanni, A., D. Folloni, L. Verhagen, J. Sallet, M.C. Klein-Flügge, and M.F. Rushworth. 2021. Activation and disruption of a neural mechanism for novel choice in monkeys. Nature 591 (7849): 270–274.

Bottini, R. and C.F. Doeller. 2020. Knowledge across reference frames: Cognitive maps and image spaces. Trends in Cognitive Sciences 24 (8): 606–619.

Botvinick, M. and M. Toussaint. 2012. Planning as inference. Trends in cognitive sciences 16 (10): 485–488.

Buesing, L., J. Bill, B. Nessler, and W. Maass. 2011. Neural dynamics as sampling: a model for stochastic computation in recurrent networks of spiking neurons. PLoS computational biology 7 (11): e1002211.

Bush, D., C. Barry, D. Manson, and N. Burgess. 2015. Using grid cells for navigation. Neuron 87 (3): 507–520.

Buzsáki, G. and E.I. Moser. 2013. Memory, navigation and theta rhythm in the hippocampal-entorhinal system. Nature neuroscience 16 (2): 130–138.

Campbell, M.G., S.A. Ocko, C.S. Mallory, I.I. Low, S. Ganguli, and L.M. Giocomo. 2018. Principles governing the integration of landmark and self-motion cues in entorhinal cortical codes for navigation. Nature neuroscience 21 (8): 1096–1106.

Chandra, S., S. Sharma, R. Chaudhuri, and I. Fiete. 2025. Episodic and associative memory from spatial scaffolds in the hippocampus. Nature 638 (8051): 739–751.

Constantinescu, A.O., J.X. O’Reilly, and T.E. Behrens. 2016. Organizing conceptual knowledge in humans with a gridlike code. Science 352 (6292): 1464–1468.

Crivelli-Decker, J., A. Clarke, S.A. Park, D.J. Huffman, E.D. Boorman, and C. Ranganath. 2023. Goal-oriented representations in the human hippocampus during planning and navigation. Nature communications 14 (1): 2946.

Davies, M., A. Wild, G. Orchard, Y. Sandamirskaya, G.A.F. Guerra, P. Joshi, P. Plank, and S.R. Risbud. 2021. Advancing neuromorphic computing with loihi: A survey of results and outlook. Proceedings of the IEEE 109 (5): 911–934.

Dong, L.L. and I.R. Fiete. 2024. Grid cells in cognition: mechanisms and function. Annual Review of Neuroscience 47.

Eppstein, D. 1998. Finding the k shortest paths. SIAM Journal on computing 28 (2): 652–673.

Epstein, R.A., E.Z. Patai, J.B. Julian, and H.J. Spiers. 2017. The cognitive map in humans: spatial navigation and beyond. Nature neuroscience 20 (11): 1504–1513.

FeldmanHall, O., J.Y. Son, and A. Bhandari. 2025. Abstract cognitive maps for complex social systems. Current Directions in Psychological Science: 09637214251342742.

Fiser, J., P. Berkes, G. Orbán, and M. Lengyel. 2010. Statistically optimal perception and learning: from behavior to neural representations. Trends in cognitive sciences 14 (3): 119–130.

Flerova, N., R. Marinescu, and R. Dechter. 2016. Searching for the m best solutions in graphical models. Journal of Artificial Intelligence Research 55: 889–952.

Foster, D.J. 2017. Replay comes of age. Annual review of neuroscience 40 (1): 581–602.

Friston, K. 2010. The free-energy principle: a unified brain theory? Nature reviews neuroscience 11 (2): 127–138.

Garcia, C.E., D.M. Prett, and M. Morari. 1989. Model predictive control: Theory and practice—a survey. Automatica 25 (3): 335–348.

Gardner, R.J., E. Hermansen, M. Pachitariu, Y. Burak, N.A. Baas, B.A. Dunn, M.B. Moser, and E.I. Moser. 2022. Toroidal topology of population activity in grid cells. Nature 602 (7895): 123–128.

Gauthier, J.L. and D.W. Tank. 2018. A dedicated population for reward coding in the hippocampus. Neuron 99 (1): 179–193.

Geffner, H. and B. Bonet. 2013. A concise introduction to models and methods for automated planning. Morgan & Claypool Publishers.

Gelly, S. and D. Silver. 2011. Monte-carlo tree search and rapid action value estimation in computer go. Artificial Intelligence 175 (11): 1856–1875.

George, D., R.V. Rikhye, N. Gothoskar, J.S. Guntupalli, A. Dedieu, and M. Lázaro-Gredilla. 2021. Clone-structured graph representations enable flexible learning and vicarious evaluation of cognitive maps. Nature communications 12 (1): 2392.

Gershman, S.J., J.A. Assad, S.R. Datta, S.W. Linderman, B.L. Sabatini, N. Uchida, and L. Wilbrecht. 2024. Explaining dopamine through prediction errors and beyond. Nature neuroscience 27 (9): 1645–1655.

Gupta, A.S., M.A. Van Der Meer, D.S. Touretzky, and A.D. Redish. 2010. Hippocampal replay is not a simple function of experience. Neuron 65 (5): 695–705.

Habenschuss, S., Z. Jonke, and W. Maass. 2013. Stochastic computations in cortical microcircuit models. PLoS computational biology 9 (11): e1003311.

Hafting, T., M. Fyhn, S. Molden, M.B. Moser, and E.I. Moser. 2005. Microstructure of a spatial map in the entorhinal cortex. Nature 436 (7052): 801–806.

Hart, P.E., N.J. Nilsson, and B. Raphael. 1968. A formal basis for the heuristic determination of minimum cost paths. IEEE transactions on Systems Science and Cybernetics 4 (2): 100–107.

Hills, T.T., P.M. Todd, D. Lazer, A.D. Redish, and I.D. Couzin. 2015. Exploration versus exploitation in space, mind, and society. Trends in cognitive sciences 19 (1): 46–54.

Hok, V., P.P. Lenck-Santini, S. Roux, E. Save, R.U. Muller, and B. Poucet. 2007. Goal-related activity in hippocampal place cells. Journal of Neuroscience 27 (3): 472–482.

Høydal, Ø.A., E.R. Skytøen, S.O. Andersson, M.B. Moser, and E.I. Moser. 2019. Object-vector coding in the medial entorhinal cortex. Nature 568 (7752): 400–404.

Iyer, A., S. Chandra, S. Sharma, and I. Fiete. 2024. Flexible mapping of abstract domains by grid cells via self-supervised extraction and projection of generalized velocity signals. Advances in Neural Information Processing Systems 37: 85441–85466.

Jeong, H., A. Taylor, J.R. Floeder, M. Lohmann, S. Mihalas, B. Wu, M. Zhou, D.A. Burke, and V.M.K. Namboodiri. 2022. Mesolimbic dopamine release conveys causal associations. Science 378 (6626): eabq6740.

Jonke, Z., S. Habenschuss, and W. Maass. 2016. Solving constraint satisfaction problems with networks of spiking neurons. Frontiers in neuroscience 10: 118.

Kahnt, T. and G. Schoenbaum. 2025. The curious case of dopaminergic prediction errors and learning associative information beyond value. Nature Reviews Neuroscience: 1–10.

Kawato, M. 1999. Internal models for motor control and trajectory planning. Current opinion in neurobiology 9 (6): 718–727.

Kobilarov, M. 2012. Cross-entropy motion planning. The International Journal of Robotics Research 31 (7): 855–871.

Kurth-Nelson, Z., T. Behrens, G. Wayne, K. Miller, L. Luettgau, R. Dolan, Y. Liu, and P. Schwartenbeck. 2023. Replay and compositional computation. Neuron 111 (4): 454–469.

Lin, H. and Y. Yang. 2026, April. LH-cbicr/GCML: GCML. 10.5281/zenodo.19370442. Zenodo, Version GCML, doi:10.5281/zenodo.19370442.

Linares López, C. and I. Herman 2024, October. Evolving a* to efficiently solve the κ shortest-path problem. In U. Endriss, F. S. Melo, K. Bach, A. Bugarín-Diz, J. M. Alonso-Moral, S. Barro, and F. Heintz (Eds.), ECAI 2024: 27th European Conference on Artificial Intelligence, 19–24 October 2024, Santiago de Compostela, Spain (Including PAIS 2024), Volume 392 of Frontiers in Artificial Intelligence and Applications, Amsterdam, The Netherlands, pp. s4352–4359. IOS Press.

Liu, C., R. Todorova, W. Tang, A. Oliva, and A. Fernandez-Ruiz. 2023. Associative and predictive hippocampal codes support memory-guided behaviors. Science 382 (6668): eadi8237.

Liu, G., M. Tang, and B. Eysenbach. 2024. A single goal is all you need: Skills and exploration emerge from contrastive rl without rewards, demonstrations, or subgoals. arXiv preprint 2408.05804

Ma, W.J., J.M. Beck, P.E. Latham, and A. Pouget. 2006. Bayesian inference with probabilistic population codes. Nature neuroscience 9 (11): 1432–1438.

Maass, W. 2014. Noise as a resource for computation and learning in networks of spiking neurons. Proceedings of the IEEE 102 (5): 860–880.

Maass, W., T. Natschläger, and H. Markram. 2002. Real-time computing without stable states: A new framework for neural computation based on perturbations. Neural computation 14 (11): 2531–2560.

Manns, J.R. and H. Eichenbaum. 2009. A cognitive map for object memory in the hippocampus. Learning & memory 16 (10): 616–624.

McNaughton, B.L., F.P. Battaglia, O. Jensen, E.I. Moser, and M.B. Moser. 2006. Path integration and the neural basis of the’cognitive map’. Nature Reviews Neuroscience 7 (8): 663–678.

Milstein, A.D., Y. Li, K.C. Bittner, C. Grienberger, I. Soltesz, J.C. Magee, and S. Romani. 2021. Bidirectional synaptic plasticity rapidly modifies hippocampal representations. Elife 10: e73046.

Moore, C. and J.M. Robson. 2001. Hard tiling problems with simple tiles. Discrete & Computational Geometry 26: 573–590.

Moser, E.I., E. Kropff, and M.B. Moser. 2008. Place cells, grid cells, and the brain’s spatial representation system. Annu. Rev. Neurosci. 31 (1): 69–89.

Muhle-Karbe, P.S., H. Sheahan, G. Pezzulo, H.J. Spiers, S. Chien, N.W. Schuck, and C. Summerfield. 2023. Goal-seeking compresses neural codes for space in the human hippocampus and orbitofrontal cortex. Neuron 111 (23): 3885–3899.

Nessler, B., M. Pfeiffer, L. Buesing, and W. Maass. 2013. Bayesian computation emerges in generic cortical microcircuits through spike-timing-dependent plasticity. PLoS computational biology 9 (4): e1003037.

Newell, A., H.A. Simon, et al. 1972. Human problem solving, Volume 104. Prentice-hall Englewood Cliffs, NJ.

Nyberg, N., É. Duvelle, C. Barry, and H.J. Spiers. 2022. Spatial goal coding in the hippocampal formation. Neuron 110 (3): 394–422.

O’Keefe, J. and L. Nadel. 1978. The hippocampus as a cognitive map. Oxford University Press, New York.

Ólafsdóttir, H.F., D. Bush, and C. Barry. 2018. The role of hippocampal replay in memory and planning. Current Biology 28 (1): R37–R50.

Park, S.A., D.S. Miller, and E.D. Boorman. 2021. Inferences on a multidimensional social hierarchy use a grid-like code. Nature neuroscience 24 (9): 1292–1301.

Parr, T., G. Pezzulo, and K.J. Friston. 2022. Active inference: the free energy principle in mind, brain, and behavior. MIT Press.

Pennartz, C., R. Ito, P. Verschure, F. Battaglia, and T. Robbins. 2011. The hippocampal–striatal axis in learning, prediction and goal-directed behavior. Trends in neurosciences 34 (10): 548–559.

Pezzulo, G., M.A. Van der Meer, C.S. Lansink, and C.M. Pennartz. 2014. Internally generated sequences in learning and executing goal-directed behavior. Trends in cognitive sciences 18 (12): 647–657.

Pfeiffer, B.E. and D.J. Foster. 2013. Hippocampal place-cell sequences depict future paths to remembered goals. Nature 497 (7447): 74–79.

Pouget, A., P. Dayan, and R.S. Zemel. 2003. Inference and computation with population codes. Annual review of neuroscience 26 (1): 381–410.

Powers, W.T. 1973. Behavior: The control of perception. Aldine Chicago.

Raju, R.V., J.S. Guntupalli, G. Zhou, C. Wendelken, M. Lázaro-Gredilla, and D. George. 2024. Space is a latent sequence: A theory of the hippocampus. Science Advances 10 (31): eadm8470.

Redish, A.D. 2016. Vicarious trial and error. Nature Reviews Neuroscience 17 (3): 147–159.

Rivard, B., Y. Li, P.P. Lenck-Santini, B. Poucet, and R.U. Muller. 2004. Representation of objects in space by two classes of hippocampal pyramidal cells. Journal of General Physiology 124 (1): 9–25.

Rosenblueth, A., N. Wiener, and J. Bigelow. 1943. Behavior, purpose and teleology. Philosophy of science 10 (1): 18–24.

Russell, S.J. and P. Norvig. 2016. Artificial intelligence: a modern approach. pearson.

Schuck, N.W., M.B. Cai, R.C. Wilson, and Y. Niv. 2016. Human orbitofrontal cortex represents a cognitive map of state space. Neuron 91 (6): 1402–1412.

Schwartenbeck, P., A. Baram, Y. Liu, S. Mark, T. Muller, R. Dolan, M. Botvinick, Z. Kurth-Nelson, and T. Behrens. 2023. Generative replay underlies compositional inference in the hippocampal-prefrontal circuit. Cell 186 (22): 4885–4897.

Sorscher, B., G.C. Mel, S.A. Ocko, L.M. Giocomo, and S. Ganguli. 2023. A unified theory for the computational and mechanistic origins of grid cells. Neuron 111 (1): 121–137.

Stachenfeld, K.L., M.M. Botvinick, and S.J. Gershman. 2017. The hippocampus as a predictive map. Nature neuroscience 20 (11): 1643–1653.

Stöckl, C., Y. Yang, and W. Maass. 2024. Local prediction-learning in high-dimensional spaces enables neural networks to plan. Nature Communications 15 (1): 2344.

Stoianov, I., D. Maisto, and G. Pezzulo. 2022. The hippocampal formation as a hierarchical generative model supporting generative replay and continual learning. Progress in Neurobiology 217: 102329.

Sutton, R.S. 1992. Adapting bias by gradient descent: An incremental version of delta-bar-delta. In AAAI, pp. 171–176. Citeseer.

Sutton, R.S. and A.G. Barto. 2018. Reinforcement learning: An introduction. MIT press.

Tavares, R.M., A. Mendelsohn, Y. Grossman, C.H. Williams, M. Shapiro, Y. Trope, and D. Schiller. 2015. A map for social navigation in the human brain. Neuron 87 (1): 231–243.

Tsoar, A., R. Nathan, Y. Bartan, A. Vyssotski, G. Dell’Omo, and N. Ulanovsky. 2011. Large-scale navigational map in a mammal. Proceedings of the National Academy of Sciences 108 (37): E718–E724.

Ujfalussy, B.B. and G. Orbán. 2022. Sampling motion trajectories during hippocampal theta sequences. Elife 11: e74058.

Ulsaker-Janke, I., T. Waaga, T. Waaga, E.I. Moser, and M.B. Moser. 2023. Grid cells in rats deprived of geometric experience during development. Proceedings of the National Academy of Sciences 120 (41): e2310820120.

Van Der Meer, M., Z. Kurth-Nelson, and A.D. Redish. 2012. Information processing in decision-making systems. The Neuroscientist 18 (4): 342–359.

Van Hasselt, H., A. Guez, and D. Silver 2016. Deep reinforcement learning with double q-learning. In Proceedings of the AAAI conference on artificial intelligence, Volume 30.

Veselic, S., T.H. Muller, E. Gutierrez, T.E. Behrens, L.T. Hunt, J.L. Butler, and S.W. Kennerley. 2025. A cognitive map for value-guided choice in the ventromedial prefrontal cortex. Cell 188 (12): 3259–3273.

Viganò, S., R. Bayramova, C.F. Doeller, and R. Bottini. 2024. Spontaneous eye movements reflect the representational geometries of conceptual spaces. Proceedings of the National Academy of Sciences 121 (17): e2403858121.

Vollan, A.Z., R.J. Gardner, M.B. Moser, and E.I. Moser. 2025. Left–right-alternating theta sweeps in entorhinal–hippocampal maps of space. Nature 639 (8056): 995–1005.

Wang, Z., T. Schaul, M. Hessel, H. Hasselt, M. Lanctot, and N. Freitas 2016. Dueling network architectures for deep reinforcement learning. In International conference on machine learning, pp. 1995–2003. PMLR.

Whittington, J.C., T.H. Muller, S. Mark, G. Chen, C. Barry, N. Burgess, and T.E. Behrens. 2020. The tolman-eichenbaum machine: Unifying space and relational memory through generalization in the hippocampal formation. Cell 183 (5): 1249–1263.

Widloski, J. and D.J. Foster. 2022. Flexible rerouting of hippocampal replay sequences around changing barriers in the absence of global place field remapping. Neuron 110 (9): 1547–1558.

Wiener, N. 1961. Cybernetics or Control and Communication in the Animal and the Machine, Volume 25. MIT press.

Wittkuhn, L., S. Chien, S. Hall-McMaster, and N.W. Schuck. 2021. Replay in minds and machines. Neuroscience & Biobehavioral Reviews 129: 367–388.

Wolpert, D.M. and M. Kawato. 1998. Multiple paired forward and inverse models for motor control. Neural networks 11 (7-8): 1317–1329.

Wu, Y. and W. Maass. 2025. A simple model for behavioral time scale synaptic plasticity (btsp) provides content addressable memory with binary synapses and one-shot learning. Nature communications 16 (1): 342.

Xiao, Z., X. Wang, J. Zhang, J. Ou, L. He, Y. Qu, X. Hu, T. Behrens, and Y. Liu. 2025. Human hippocampal ripples predict the alignment of experience to a grid-like schema. bioRxiv : 2025–01.

Yang, Y. and W. Maass. 2025. Episodic memories make goal directed action selection context-aware and explainable. bioRxiv : 2025–10.

Zutshi, I., A. Apostolelli, W. Yang, Z.S. Zheng, T. Dohi, E. Balzani, A.H. Williams, C. Savin, and G. Buzsáki. 2025. Hippocampal neuronal activity is aligned with action plans. Nature: 1–9.

